# Spatiotemporal regulation of ciliary outer dynein arm biogenesis and the role of PIH homologs

**DOI:** 10.1101/2025.07.26.666928

**Authors:** Karthika Balasubramaniam, Fangqi Fu, Cynthia Y. He

**Affiliations:** Department of Biological Sciences National University of Singapore 14 Science Drive 4, Singapore 117543

## Abstract

Outer Dynein Arm complex (ODA) powers motile cilia that are indispensable for fundamental processes, including embryogenesis, reproduction, and respiration. The ODA subunits: heavy (HC), intermediate (IC), and light chains (LC), are known to preassemble in the cytoplasm with the help of Dynein Axonemal Assembly Factors (DNAAFs). The mechanistic details underlying functional ODA assembly and the specific roles of DNAAFs in orchestrating subunit translation, folding, and assembly are still poorly defined. Using *Trypanosoma brucei* as a model, we showed that ODA preassembly is coupled to HC translation, during which ICs post-translationally associate with translating HCs to form co-translational assembly sites. PIH1D1, a DNAAF and a PIH family protein, concentrates at these cotranslational assembly sites. Without PIH1D1, HC protein levels are reduced, leaving the IC-LC complex stranded in the cytoplasm. We propose that PIH1D1 generates specialised compartments to assist the co-translational folding of HCs and enable their assembly with the other ODA subunits. Our study highlights spatial compartmentalization as a conserved strategy that interweaves translation, folding, and assembly to ensure ordered and timely ODA formation during ciliogenesis.

## Introduction

Assembling proteins into higher-order complexes with precise spatial-temporal control is critical for fundamental functions ranging from gene regulation to cell division to structural integrity. However, achieving such assembly within a crowded and dynamic cellular milieu poses a considerable challenge. Cells employ various mechanisms to assemble protein complexes, with the two well-known being post- and co-translational assembly. In post-translational assembly, the proteins are synthesised individually before assembling into complexes. In contrast, in cotranslational assembly, proteins assemble with their partners as soon as the interaction domains emerge from the translating ribosomes (Morales-Polanco et al., 2022; Schwarz & Beck, 2019). Although post-translational assembly is viewed as the predominant mechanism, growing evidence indicates that the co-translational assembly is pervasive across eukaryotes (Bertolini et al., 2021; Duncan & Mata, 2011; Kamenova et al., 2019; Shiber et al., 2018). Nonetheless, the molecular basis of what dictates the choice of a particular assembly pathway and the regulatory mechanisms involved for a given protein complex is poorly defined.

Here, we investigated the assembly of the Outer Dynein Arm (ODA), an evolutionarily conserved motor complex crucial for ciliary motility. This multi-subunit, mega-dalton complex typically comprises two to three heavy chains (HCs), two intermediate chains (ICs), and a group of light chains (LCs) (King, 2016). Although ODA functions within cilia, its subunits are synthesised and assembled step by step in the cytoplasm before being shipped to their site of action in cilia (Fowkes & Mitchell, 1998). The successful packaging of ODA subunits into a mature complex relies on the concerted action of a group of cytoplasmic factors named Dynein Axonemal Assembly Factors or DNAAFs (Braschi et al., 2022; Desai et al., 2018). Any disruption in this process due to defective ODA subunits or DNAAFs compromises ciliary motility, resulting in a wide range of human diseases collectively known as motile ciliopathies (Braschi et al., 2022; Wallmeier et al., 2020; Zariwala et al., 1993). Despite the identification of DNAAFs, their specific roles in facilitating the ordered and timely assembly of the ODA during ciliogenesis are not fully understood.

In this study, we focused on a subset of DNAAFs that belong to the PIH protein family, all of which contain a conserved PIH1 domain (Proteins Interacting with Hsp90 1) (Zhao et al., 2005). The PIH1 protein, together with RuvB DNA helicases 1 & 2 and a TPR domain-containing protein Tah1, form the R2TP complex, a well-studied Hsp90 co-chaperone (Kakihara & Houry, 2012). In eukaryotes, R2TP has emerged as a key player in assembling various multisubunit protein complexes, including RNA polymerase II (Boulon et al., 2010), small ribonuclear protein (snoRNP) (Kakihara et al., 2014; Zhao et al., 2008), and PIKK signaling-related protein complexes (Horejsi et al., 2010; Izumi et al., 2010; Pal et al., 2014). Several studies have also reported the participation of the R2TP complex in assembling ciliary dyneins (Dong et al., 2014; Y. Li et al., 2017; Olcese et al., 2017; Omran et al., 2008; Paff et al., 2017). Intriguingly, organisms with motile cilia contain three or four distinct PIH proteins, as compared to organisms lacking motile cilia that contain one or none (Yamamoto et al., 2010). Given that the PIH component is involved in client recognition (Horejsi et al., 2014), the presence of multiple PIH homologs allows R2TP to earmark distinct types of ciliary dyneins and target them to molecular chaperones for proper folding and assembly. Consistent with this idea, functional analyses in *Zebrafish* and *Chlamydomonas* (Yamaguchi et al., 2018; Yamamoto et al., 2020) highlighted the essential role of PIH homologs in aiding the assembly of diverse types of ciliary dyneins, such as the ODA and the inner dynein arm (IDA).

*Trypanosoma brucei* is a unicellular eukaryote with highly conserved motile ciliary machinery. In our recent work, we demonstrated the conservation of DNAAFs and a stepwise cytoplasmic preassembly pathway for the ODA, similar to other eukaryotes (Balasubramaniam et al., 2024). In this study, we show that PIH1D1, one of the four PIH homologs in *Trypanosoma brucei,* is essential for the ODA assembly. Importantly, during ciliogenesis, PIH1D1 provides a unique niche to support the folding of nascent heavy chains and their co-translational assembly with ICs and LCs. Our findings demonstrated that ODA subunits assemble in a defined order that is dictated by the localized translation of the HCs, a principle likely conserved in all eukaryotes containing motile cilia.

## Results

### 1. *In situ* proximity mapping of PIH homologs

*T. brucei* genome encodes four distinct PIH homologs, which are named KTU (Tb927.10.12860), PIH1D1 (Tb927.9.10490), PIH1D2 (Tb927.10.16020), and PIH1D3 (Tb927.3.4410) to partially reflect their phylogenetic grouping with other PIH proteins (Fig. S1A). PIH1D1, also known as CMF56, is essential for the motility of the parasite (Baron et al., 2007). All PIH homologs, when depleted by RNAi, impaired parasite motility and growth (Fig. S1B). Despite the conserved role in ciliary dynein assembly, the PIH homologs display poor conservation in dynein client specificity across species (Olcese et al., 2017; Paff et al., 2017; Yamaguchi et al., 2018; Yamamoto et al., 2020). To establish the specific roles of each PIH protein in ciliary dynein assembly, we utilized BioID2 (Kim et al., 2016) to map their proximity interactors. Each PIH homolog was tagged with 3HA-BioID2 at its N-terminus and expressed using a cumate inducible system (F. J. Li et al., 2017). Upon biotinylation, the proximal partners of each PIH homolog were affinity-purified and identified by LC-MS/MS.

A total of 18, 44, 37, and 99 proteins were identified using KTU, PIH1D1, PIH1D2, and PIH1D3 as baits, respectively (Fig. 1A, Supplementary Data 1). The BioID candidates were grouped into four functional categories: 1. chaperone-related; 2. R2TP, dynein components, and DNAAF’s; 3. Translation-related and 4. mRNA/RNA binding and stress granular components (Fig.1B). The presence of R2TP components in all interactomes suggested that all PIH homologs function as a conserved component of the R2TP complex. Their interaction with various molecular chaperones, including Hsp90, Hsp70, prefoldin, and T-complex proteins, further supported their functions as co-chaperones. Two other DNAAFs, DYX1C1 (DNAAF4) and WDR92 (DNAAF10), were found in KTU- and PIH1D1-BioID. This is consistent with a previous report showing DYX1C1 interaction with KTU in HEK293 cells, and mutations in DYX1C1 disrupt ODA and IDA assembly in human PCD patients (Tarkar et al., 2013). WDR92, although not linked to human PCD, supports dynein assembly in motile organisms such as *Chlamydomonas* (Liu et al., 2019; Patel-King et al., 2019) and *Drosophila* (Zur Lage et al., 2018). Notably, among the PIH homologs, only PIH1D1 BioID specifically detected ODAα, a heavy chain component of ODA, as a high-confidence hit. The absence of known ciliary clients in the interactomes of other PIH proteins may be due to detection limits or novel clients that are yet to be characterized.

**Figure 1.**
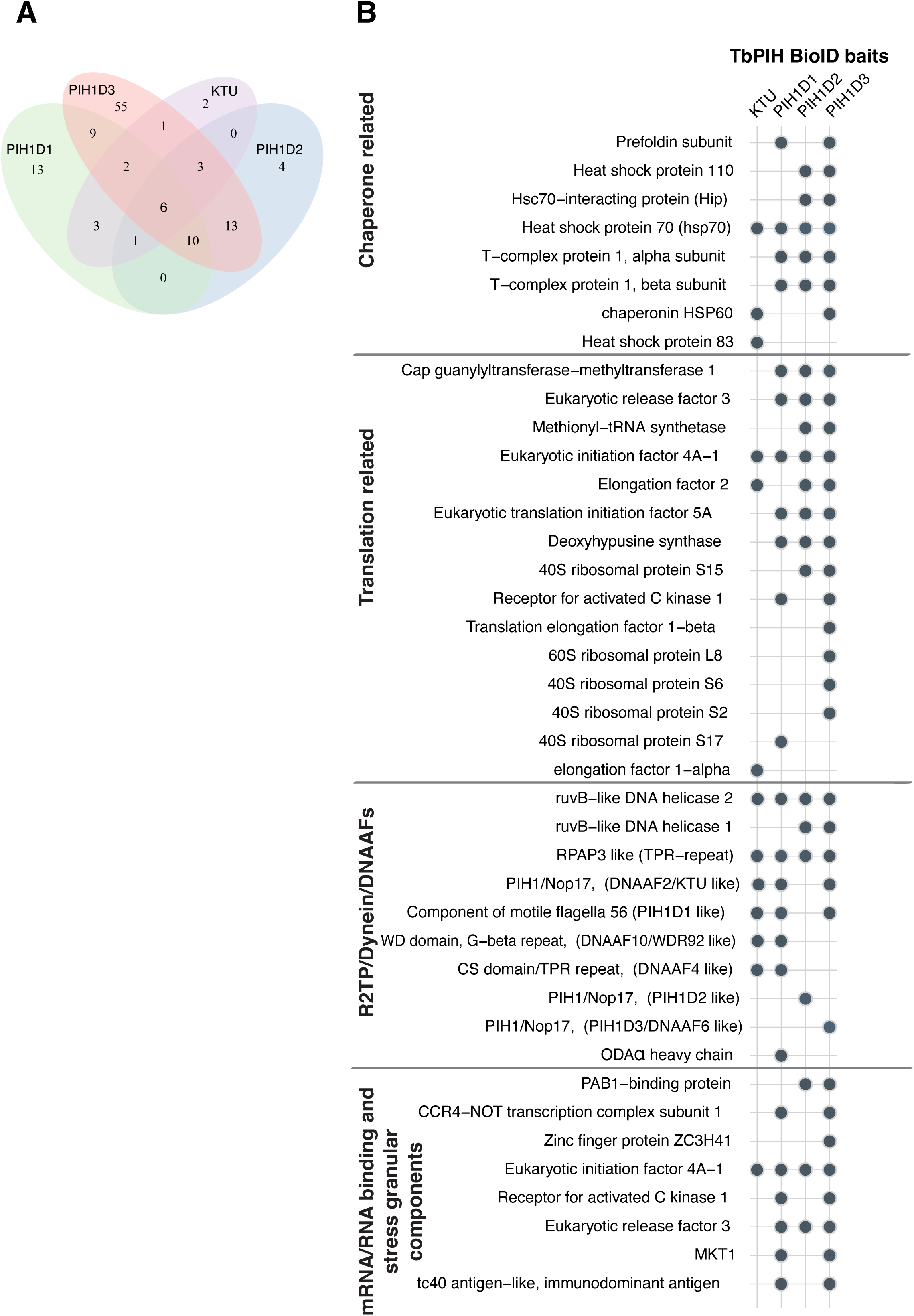
BioID interactome of PIH homologs in *Trypanosoma brucei*. **A)** Venn diagram showing the unique and overlapping high-confidence (based on emPAI score) interactors among the PIH homologs. **B)** Dot plot illustration of the selected proteins from four major functional categories identified in the BioID interactome of all four PIH homologs. Each dot represents the presence of a protein in the specified interactome.

All PIH homologs were additionally associated with core translational machinery components, including EIF4A-1, EIF5A, and ERF3 (Fig. 1B), suggesting a potential role in protein translation. Moreover, several mRNA/RNA-binding proteins, such as CGM1, MKT1, CNOT1, and PBP1, involved in various aspects of RNA metabolism, were also present. Some of these proteins, such as CNOT1, MKT1, and PBP1 (ataxin2 in mammals), are components of stress granules or P-bodies in yeast (Lewis et al., 2014; Muhlrad & Parker, 2005; Swisher & Parker, 2010), mammals (Nonhoff et al., 2007; Tsai et al., 2025), and *T. brucei* (Fritz et al., 2015; Singh et al., 2014). Together, these data suggest that PIH homologs, beyond their role in dynein assembly, may also be involved in protein translation and RNA metabolism.

### 2. PIH1D1 is required for the ODA preassembly in *T. brucei*

As ODAα was identified in PIH1D1 BioID, we hypothesized that PIH1D1 plays a role in ODA assembly. To test this, we generated ODA reporter cell lines by endogenously tagging the core subunits of the ODA – ODAα and ODAβ, IC1 and IC2 – with Ty:mNG:Ty at their N-terminal ends, as described in our previous study (Balasubramaniam et al., 2024). All tagged ODA reporters displayed a strong signal along the flagellum, overlapping with an axonemal marker mAb25 (Dacheux et al., 2012) in both the old and new flagellum in duplicating cells (Fig. 2A & C). Upon PIH1D1 RNAi, there was a marked reduction in ciliary incorporation of all tested ODA subunits (Fig. 2A, B, C & D, Fig. S2A & B). But unlike the heavy chains, the intermediate chains appeared to accumulate in the cytosol in PIH1D1 RNAi cells. To investigate this differential accumulation of ODA subunits in PIH1D1 RNAi cells, we extracted the cells using 1% NP-40 (in PEME buffer). In the control cells, both ODAα and IC1 were enriched in the detergent-insoluble, cytoskeletal fraction (Fig. 2E & F), consistent with their stable integration into the flagellar axoneme. In PIH1D1 RNAi cells, the total protein level of IC1 remained unchanged, but was reduced in the cytoskeletal pool and enriched in the cytoplasmic pool (Fig. 2F). In contrast, there was an overall reduction in the ODAα protein level across all fractions (Fig. 2E). qRT-PCR showed that the mRNA abundance of ODA components was not significantly changed following PIH1D1 RNAi (Fig. 2G), suggesting that PIH1D1 regulates the ODAα at the translational or post-translational level.

**Figure 2.**
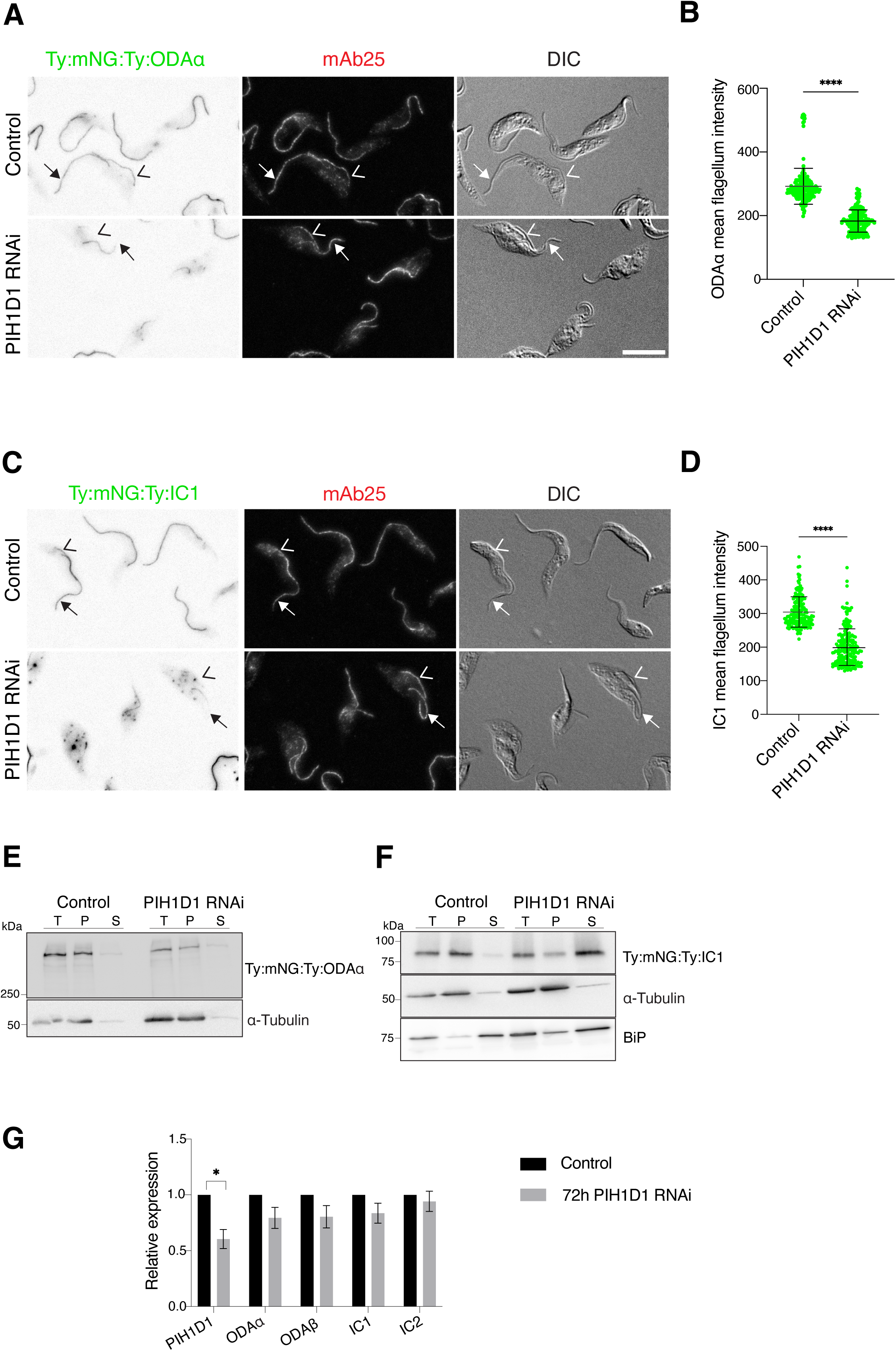
Depletion of PIH1D1 reduces the protein abundance of ODA HCs, but not the ICs. **A) & C)** Cells endogenously expressing ODAα or IC1 tagged N terminally with Ty:mNG:Ty were induced for PIH1D1 RNAi for 48h or not. The cells were additionally stained with mAb25 to label the axoneme. Arrowheads and Arrows indicate the new and old flagella, respectively. Scale bar = 5 µm **B) & D)** Quantification of mean flagellar intensity of ODAα or IC1 (as reported by mNG), measured from the distal tip of the flagellum (∼ 1 µm) using ImageJ. The mean values are from three independent biological replicates. At least 150 cells were measured for each replicate. Unpaired Two-tailed t test with Welch’s correction was performed, p<0.0001 (****). **(E-F)** Representative Western blot analysis of subcellular fractions of control or PIH1D1 RNAi (72h) cells expressing Ty:mNG:Ty:ODAα or Ty:mNG:Ty:IC1 using anti-Ty antibody. T-Total cell lysate, P-Pellet, flagellar fraction, S-Soluble, cytoplasmic fraction. **G)** qRT-PCR analysis of different ODA subunits upon PIH1D1 RNAi for 72h. The TERT gene was used as a reference. Three independent biological replicates were performed, each with three technical repeats. Unpaired Two-tailed t test with Welch’s correction was performed, p=0.0150.

To further investigate the role of PIH1D1 in ODA cytoplasmic preassembly, we prepared cytoplasmic extracts from the control and PIH1D1 RNAi cells expressing natively tagged Ty:YFP:Ty:IC1 and performed sucrose gradient ultracentrifugation. As shown in our previous study (Balasubramaniam et al., 2024), in control cells, IC1 exists in at least two different populations: a higher density population containing ODAα (fractions 17-19), likely a full ODA complex; and a lower density population (fractions 9-11), representing an intermediate complex. Upon PIH1D1 RNAi, the IC1 signal from the higher density fractions (17-19) was significantly reduced, along with ODAα. However, the IC1 signal from the lower density fractions (9-11) slightly increased (Fig. 3A).

**Figure 3.**
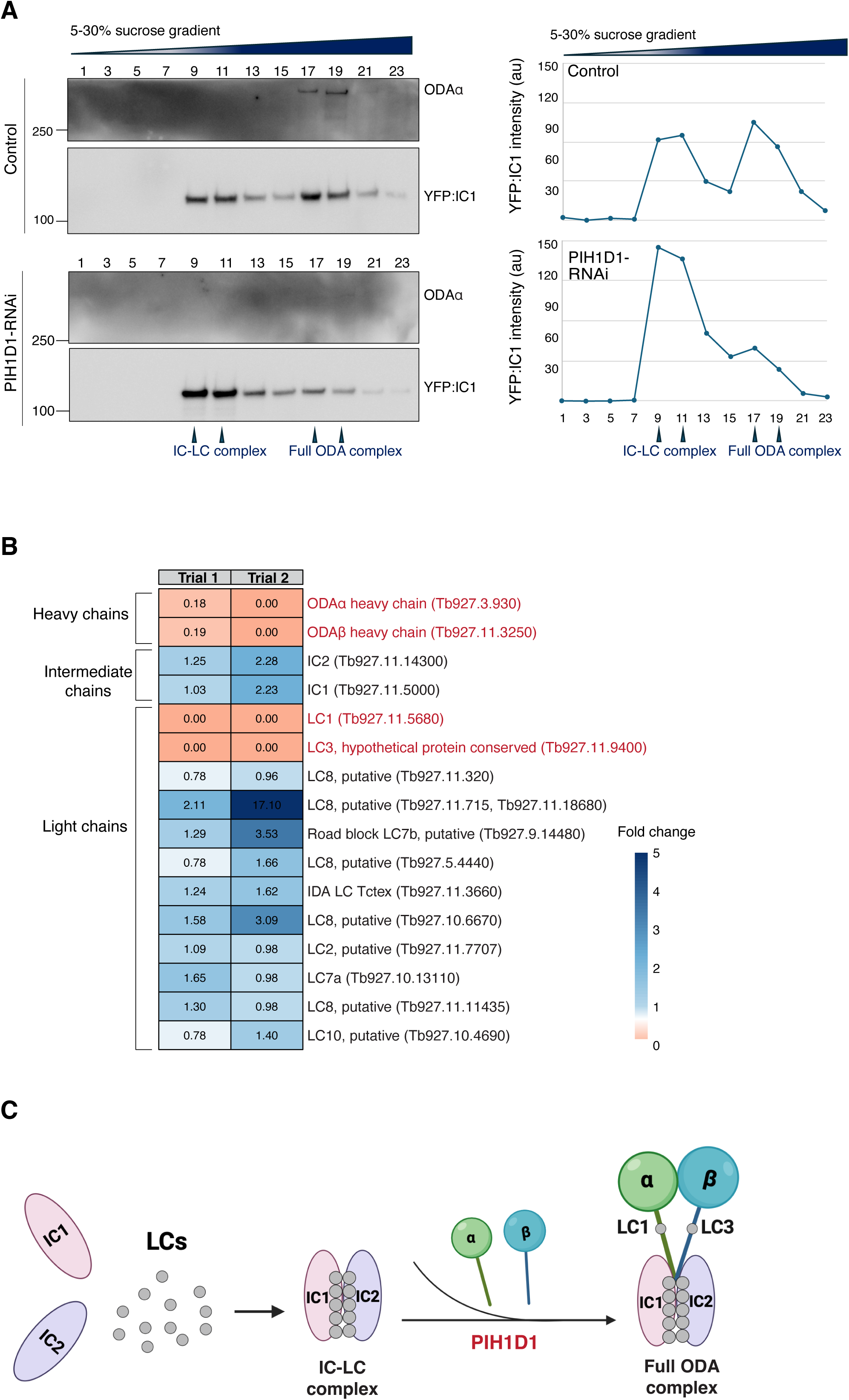
PIH1D1 is required for the stable integration of the IC-LC complex with the HCs. **A)** Cytoplasmic extracts from control and PIH1D1 RNAi cells (72h) with endogenous expression of Ty:YFP:Ty:IC1 were analyzed by sucrose gradient ultracentrifugation. The presence of Ty:YFP:Ty:IC1 and ODAα was analysed by immunoblots using anti-Ty (for Ty:YFP:Ty:IC1) and custom-made anti-ODAα antibodies. The band intensity of Ty:YFP:Ty:IC1 was measured across different fractions from both control and PIH1D1 RNAi samples and plotted as a graph. **B)** Using Ty:YFP:Ty:IC1 as a bait, ODA complexes were affinity-purified from the cytosolic extracts of the control and PIH1D1 RNAi cells after 72 h of induction. The purified samples were then analysed by LC-MS/MS. Fold change for each ODA component was calculated based on protein abundance in PIH1D1 RNAi relative to the control, and presented as a heat map. Results shown are from two independent experiments, Trial 1 and Trial 2. **C)** A model diagram illustrating the point of function of PIH1D1 in the stepwise assembly of ODA (Created in BioRender. Balasubramaniam, K. (2025) https://BioRender.com/4s75e3y).

To further assess the assembly state of ODA and identify potential regulators, we affinity-purified ODA from the cytoplasmic extracts of control and PIH1D1 RNAi cells using Ty:YFP:Ty:IC1 as a bait, followed by mass spectrometry. We performed two independent affinity purification trials, Trial 1 with cycloheximide (CHX) added in the lysis buffer and Trial 2 without. In both trials, known ODA subunits-ODAα, ODAβ, IC1, IC2, and several light chains were co-purified with IC1 in control cells. However, in PIH1D1 RNAi cells, IC1 showed reduced or no interaction with ODAα and ODAβ, as well as two light chains, LC1 and a putative LC3. IC1 association with other light chains was not affected (Fig. 3B). Together, our results suggested that PIH1D1 functions at a step where it is required for the incorporation of the ODA heavy chains with a stable IC-LC intermediate complex (Fig. 3C).

It is worth noting that YFP-IC1 co-purified several other axonemal and cytoplasmic proteins that are not recognized as components of the ODA complex or factors involved in its preassembly (Fig. S3 and Supplementary Data 2). Some of these interactions were disrupted by PIH1D1 RNAi (Fig. S3). Notably, PIH1D1 RNAi consistently perturbed the association of IC1 with an ankyrin repeat-containing protein (encoded by Tb927.4.1170) and 1,2-Dihydroxy-3-keto-5-methylthiopentene dioxygenase (encoded by Tb927.4.360) in both trials. Both proteins are localized to the axoneme with unknown functions (Billington et al., 2023; Shanmugasundram et al., 2023). Interestingly, several proteins with GO term assignment to “protein folding” and “cytoplasmic translation”, such as the HSP90 family proteins and several ribosomal proteins (Fig. S3), were selectively enriched in PIH1D1-RNAi cells, suggesting a specific association of these proteins with IC1 when PIH1D1 is absent. We also noted trial-specific differences. In trial 2, the lack of CHX in the lysis buffer led to an enrichment of cell cycle regulators such as the anaphase-promoting complex and associated proteins (Fig. S3). The significance of this observation is yet to be understood.

### 3. PIH1D1 is enriched at the translational sites of ODA heavy chains

The effect of PIH1D1 on ODAα and its association with translation-related proteins in the BioID prompted us to hypothesize that PIH1D1 may play a role in ODAα translation. We first examined the cellular localization of PIH1D1, using a custom-produced polyclonal antibody against PIH1D1 (Fig. 4A, Fig. S4A). PIH1D1 was present in the cytoplasm, with enrichment in discrete cytoplasmic foci (Fig. 4A). This staining is similar to the pattern shown in Tryptag, where a native allele of PIH1D1 was tagged and visualized with mNG reporter (Billington et al., 2023). Interestingly, not all cells exhibited PIH1D1 foci (Fig. 4A). Co-staining of basal bodies and the flagella (Fig. S4B) indicated that PIH1D1 foci are more abundant in cells with new flagellum biogenesis (Fig. 4B and Fig. S4B).

**Figure 4.**
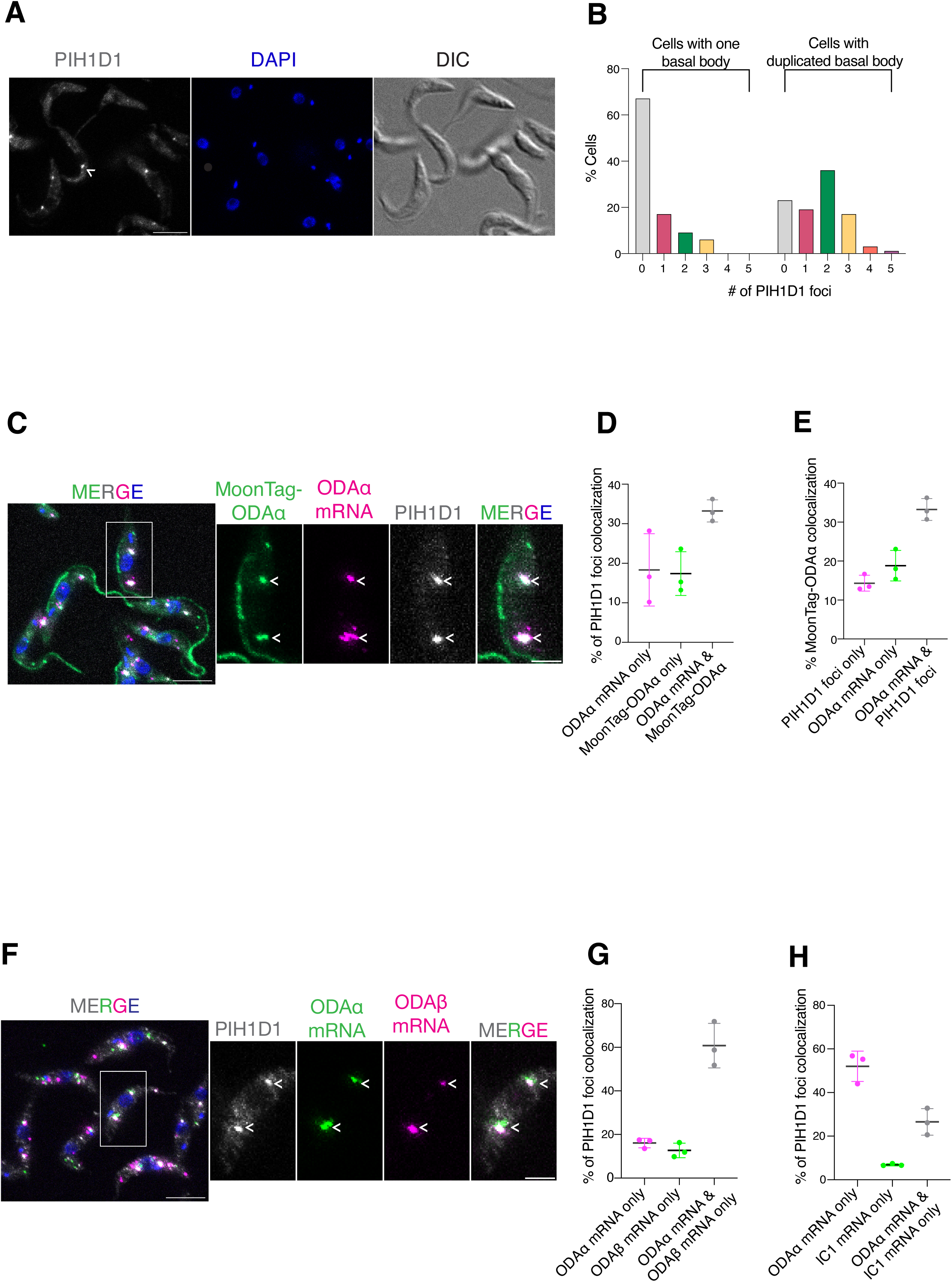
PIH1D1 is enriched at the translational sites of ODA HCs. **A)** 29.13 cells were fixed *in situ* with 4% PFA, permeabilized with 0.25% Triton X-100, and then stained with custom-made polyclonal PIH1D1 antibody and DAPI. Scale bar = 5 µm. Arrows indicate PIH1D1 foci. **B)** Quantitation plot showing the distribution of the number of PIH1D1 foci across different cell cycle stages. The different cell cycle stages were assessed by tracking the duplication of the basal body using YL1/2 antibody (see Fig. S4B). **C)** Cells expressing MoonTag-ODAα translational reporter from an endogenous allele were co-stained for ODAα mRNA and PIH1D1. Arrowheads indicate the colocalization of PIH1D1 foci with ODAα translational sites. Scale bar, Large image = 5 µm; Inset image = 2 µm. **D)** Quantitation of % of PIH1D1 foci colocalizing with ODAα mRNA only, ODAα protein only, and both. **E)** Quantitation of % MoonTag-ODAα colocalizing with PIH1D1 foci only, ODAα mRNA only, and both. Three independent biological replicates were performed. At least 250 cells were analysed for each replicate. **F)** Representative image of cells triple labelled for PIH1D1, ODAα mRNA, and ODAβ mRNA. Arrowheads indicate the colocalization of PIH1D1 foci with ODAα mRNA and ODAβ mRNA. Scale bar, Large image = 5 µm; Inset image = 2 µm. **G)** Quantitation of % of PIH1D1 foci colocalizing with ODAα mRNA, ODAβ mRNA, and both. **H)** Quantitation of % of PIH1D1 foci colocalizing with ODAα mRNA, IC1 mRNA, and both. In both **G) and H)**, the results shown were from three independent experiments with at least 500 cells quantitated in each experiment.

To visualize ODAα translation, we adopted the MoonTag system (Boersma et al., 2019). Briefly, 24 copies of the MoonTag (MT) epitope were inserted at the N-terminus of a native ODAα allele, allowing MoonTag-ODAα fusion to be expressed in cells with constitutive expression of MoonTag Binding Protein fused to sfGFP (MTBP-sfGFP). The cells expressing MTBP-sfGFP alone displayed a diffuse cytoplasmic signal (Fig. S4C), while the co-expression of MTBP-sfGFP and MoonTag-ODAα led to enrichment of the sfGFP signal along the flagellum (Fig. 4C), indicating successful incorporation of MoonTag-ODAα into the axoneme. Additionally, MoonTag-ODAα localized to cytoplasmic foci of varying size and intensity in some cells. About 50% of MoonTag-ODAα foci co-localized with ODAα mRNA, representing ODAα translational sites (Fig. 4C & E). The remaining foci not associated with ODAα mRNA may represent fully translated MoonTag-ODAα that has been released from the ribosomes.

By triple labeling MoonTag-ODAα, ODAα mRNA, and PIH1D1, we found that nearly 60 percent of MoonTag-ODAα translational sites are associated with PIH1D1 foci. On further analysis of the PIH1D1 foci, we observed four 4 distinct populations (Fig. 4D & E). ∼33 % PIH1D1 foci contained translating MoonTag-ODAα. ∼18 % PIH1D1 foci contained ODAα mRNA only, suggesting PIH1D1 association with ODAα mRNA before or soon after translation initiation. Some of these foci may also represent PIH1D1 association with untagged ODAα. ∼17 % contained MoonTag-ODAα only, suggesting PIH1D1 association with nascent MoonTag-ODAα even after it exited active translation. Finally, ∼31 % with none, PIH1D1 foci contained neither MoonTag-ODAα nor ODAα mRNA, hinting at additional roles of PIH1D1 unrelated to ODA heavy chain biogenesis.

Further, we examined the presence of transcripts from other ODA subunits, including ODAβ and IC1. Over 50% of PIH1D1 foci contained both ODAα and ODAβ mRNA (Fig. 4F & G) while PIH1D1 foci were less frequently associated with the IC1 mRNA (Fig. 4H). Together, these findings suggest that the ODA heavy chains are translated within PIH1D1 foci, while the IC1 protein is synthesized elsewhere in the cell.

### 4. The effective formation of PIH1D1 foci depends on the ODA subunits

To examine how PIH1D1 is enriched at the translational sites of ODA heavy chains and whether it depends on the expression of the ODA components per se, we knocked down ODAα, ODAβ and IC1 to evaluate their effects on PIH1D1 foci formation. In control cells, PIH1D1 displayed sharp punctate localization, whereas in RNAi cells of all tested ODA subunits, some of the PIH1D1 foci persisted, possibly due to incomplete RNAi (Fig. 5A). To eliminate any residual translation, we treated both control and RNAi cells with puromycin (Azzam & Algranati, 1973) for 30 min, which resulted in the dissipation of PIH1D1 foci in both control and RNAi cells (Fig. 5A), supporting that the formation and/or maintenance of PIH1D1 foci requires active translation. We then washed out puromycin and allowed the cells to recover for 3 h. Following the recovery, the PIH1D1 foci reappeared in the control cells, indicating the resumption of translation (Fig. 5A & C). Interestingly, RNAi silencing of the ODA subunits ODAα, ODAβ & IC1 significantly impaired the formation of PIH1D1 foci. As a control, depletion of IC138, an IDA subunit, did not perturb PIH1D1 foci formation (Fig. 5B & C).

**Figure 5.**
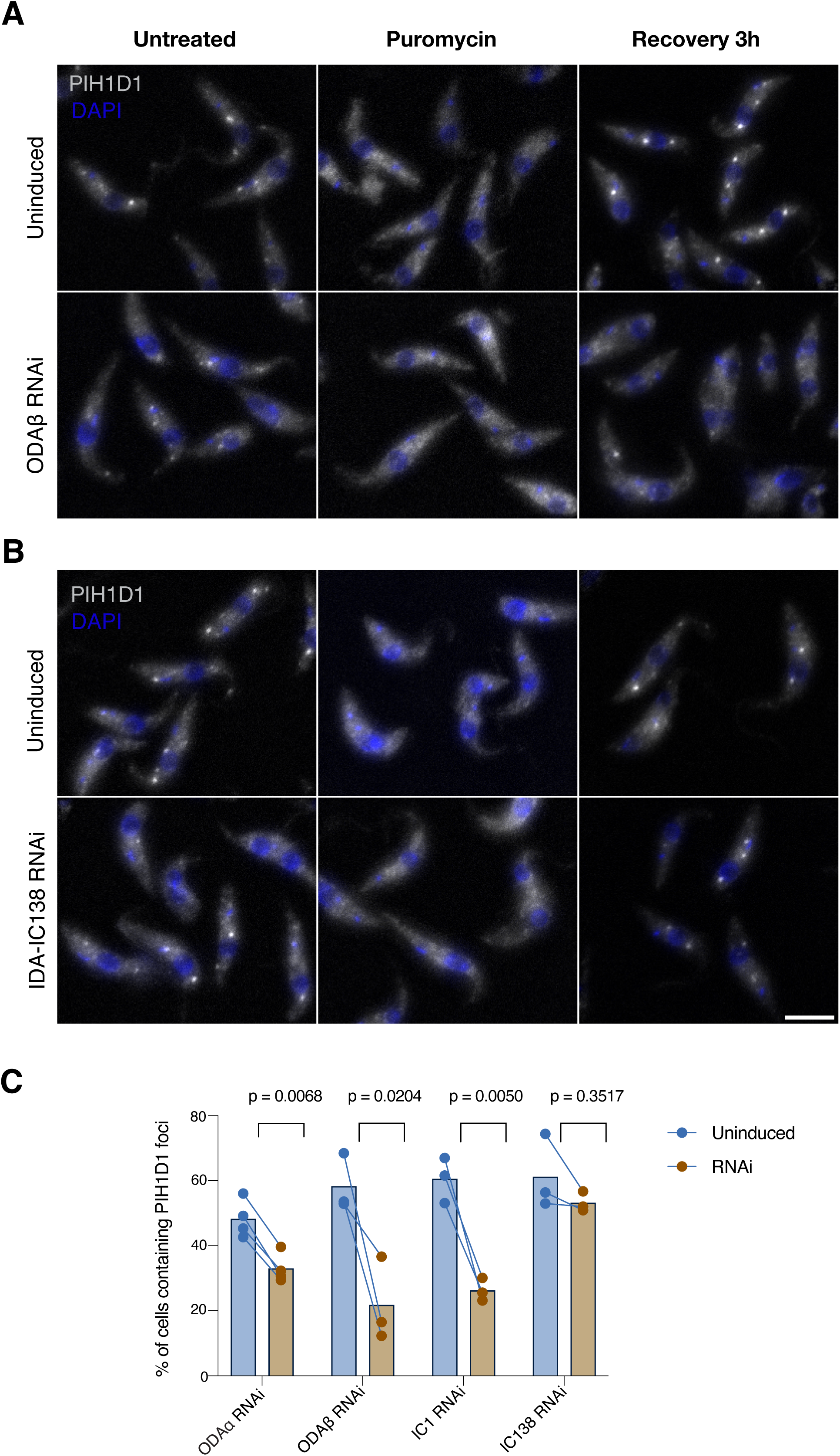
Formation of PIH1D1 foci depends on the ODA subunits. Cells endogenously expressing Ty:BioID2:Ty:PIH1D1 were treated with puromycin for 30 min. Following the puromycin treatment, the cells were washed extensively to remove puromycin and allowed to recover for 3 h. The untreated, puromycin-treated, and recovered cells were fixed *in situ* and probed with anti-Ty antibody and DAPI to visualize PIH1D1 and DNA, respectively. **A)** Cells were induced for RNAi of ODA subunits ODAα, ODAβ, and IC1, for 48h or not. A representative set of images for ODAβ RNAi is shown. **B)** As a control, cells were induced for IDA subunit IC138 RNAi for 48 h or not. **C)** Quantitation profile for the % of cells containing PIH1D1 foci upon recovery under different RNAi conditions. The lines represent the number of biological replicates. At least 150 cells were analysed for each experiment. Unpaired Two-tailed t test was performed with Welch’s correction. The corresponding *p* values are shown within the plot. Scale bar = 5 µm.

### 5. Dissecting the role of PIH1D1 in ODA heavy chain biogenesis

Our results revealed that approximately 60% of PIH1D1 foci contained both ODAα and ODAβ mRNA (Fig. 4G), suggesting translation of both heavy chains in the PIH1D1 foci. Upon puromycin treatment, the colocalization of ODAα and ODAβ mRNA was significantly reduced compared to control cells, suggesting a cotranslational interaction between the two heavy chains (Fig. 6A). Interestingly, PIH1D1 is not required for the colocalization of the two heavy chain mRNA (Fig. 6B). To investigate if PIH1D1 might be required for translation of the HCs, PIH1D1 RNAi was induced in cells stably expressing MoonTag-ODAα (Fig. 6C). Only a moderate reduction in the percentage of cells containing MoonTag-ODAα translational sites or the number of translational sites per cell was observed in PIH1D1 RNAi cells compared to control (Fig. 6D & E), suggesting that PIH1D1 is not critical for initiating MoonTag-ODAα translation.

**Figure 6.**
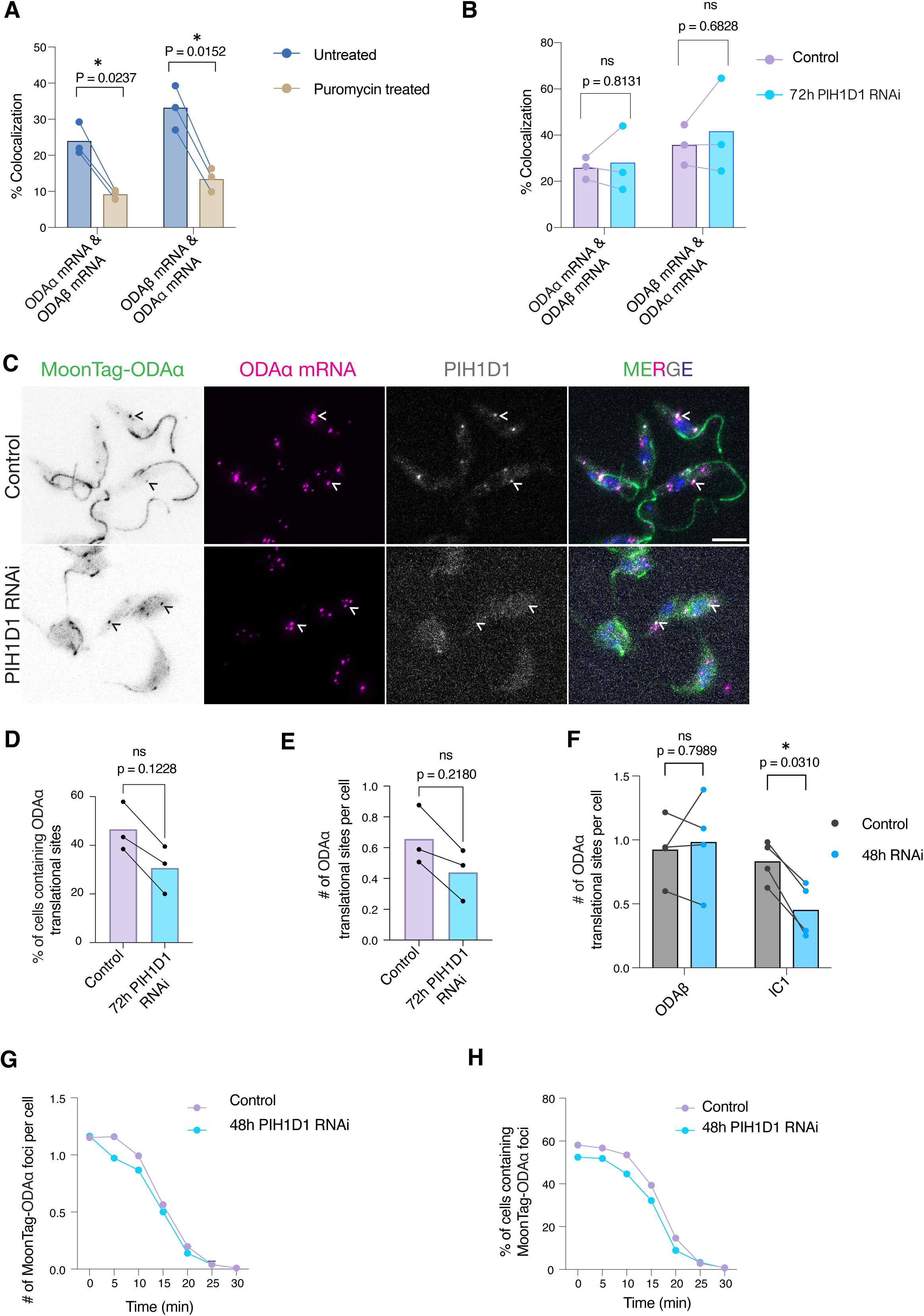
Dissecting the role of PIH1D1 in ODA heavy chain biogenesis. **(A-B)** Quantitation of the % of colocalization between ODAα mRNA and ODAβ mRNA or ODAβ mRNA and ODAα mRNA upon puromycin treatment (**A**) or PIH1D1 RNAi (72 h) (**B**). Three independent biological replicates (lines) were performed. At least 500 cells were quantified for each replicate. Unpaired Two-tailed t test with Welch’s correction was performed. **C)** Cells endogenously expressing MoonTag**-**ODAα translational reporter were induced for PIH1D1 RNAi (72h) or not, and probed for ODAα mRNA and PIH1D1. Arrowheads indicate the translational sites of ODAα colocalizing with PIH1D1 foci or not. Scale bar = 5 µm. **(D-E)** Based on the representative images shown in **C,** % of cells containing ODAα translational sites or the number of ODAα translational sites per cell was quantitated in cells with or without PIH1D1 RNAi for 72h. Three independent biological replicates (indicated by lines) were performed. At least 250 cells were quantitated for each replicate. Unpaired Two-tailed t test with Welch’s correction was performed. **F)** Cells treated with puromycin (30 min), washed and recovered for 3 or 5 h were used to quantitate the number of ODAα translational sites per cell with or without ODAβ or IC1 RNAi for 48h. Four independent biological replicates (indicated by lines) were performed. At least 140 cells per biological replicate were analysed. **(G-H)** Time course profile of harringtonine-treated cells endogenously expressing MoonTag**-**ODAα translational reporter. Harringtonine-treated control and PIH1D1 RNAi (48 h) cells were collected at specified time points and fixed *in situ* with 4 % PFA. Each data point represents the mean value from two independent biological experiments. At least 150 cells were analysed for each time point.

While the translational sites of the MoonTag-ODAα persisted in the absence of PIH1D1, those sites could contain stalled ribosomes due to errors in elongation. To examine this possibility, we performed a ribosome runoff assay using Harringtonine, which hampers translation initiation while allowing the existing ribosomes to continue translation (Fresno et al., 1977; Ingolia et al., 2011; Yan et al., 2016). Following the Harringtonine treatment, the MoonTag-ODAα foci were quantified every five minutes for 30 minutes. In control cells, the MoonTag-ODAα foci steadily dissipated over 30 minutes, indicating the completion of MoonTag-ODAα translation. In PIH1D1 RNAi cells, the dissipation kinetics of MoonTag-ODAα foci were similar to that of the control cells, suggesting that those foci are not translationally stalled products (Fig. 6G & H). Taken together, these results imply that the lack of PIH1D1 does not impede the translation initiation nor the full synthesis of the ODAα heavy chain. However, without PIH1D1, nascent ODAα protein may not be folded or assembled into the complex properly and thus is targeted for degradation soon after synthesis.

### 6. Spatio-temporal assembly of ODA complex and the role of PIH1D1 foci

While the ICs are not translated within the PIH1D1 foci, they are necessary to establish these foci. This led us to investigate if the ICs are post-translationally recruited to the PIH1D1 foci to assemble with the translating HCs. To test this, cells stably expressing Ty:mNG:Ty:IC1 were co-labelled for PIH1D1, ODAα mRNA, ODAβ mRNA, or IC1 mRNA. The results showed that IC1 protein is enriched in cytoplasmic foci that contain both PIH1D1 and ODAα mRNA (Fig. 7A). Similar to PIH1D1 foci (Fig. 4G and 4H), more than 50 percent of IC1 foci contained both ODAα and ODAβ mRNA, and less than 30 percent contained ODAα and IC1 mRNA (Fig. 7B), supporting that IC1 is post-translationally concentrated at the heavy chain translation sites. Notably, PIH1D1 RNAi did not affect the association of IC1 protein with ODAα and ODAβ mRNA (Fig. 7C, D & E), suggesting that PIH1D1 functions downstream of IC1 recruitment to HC cotranslational sites.

**Figure 7.**
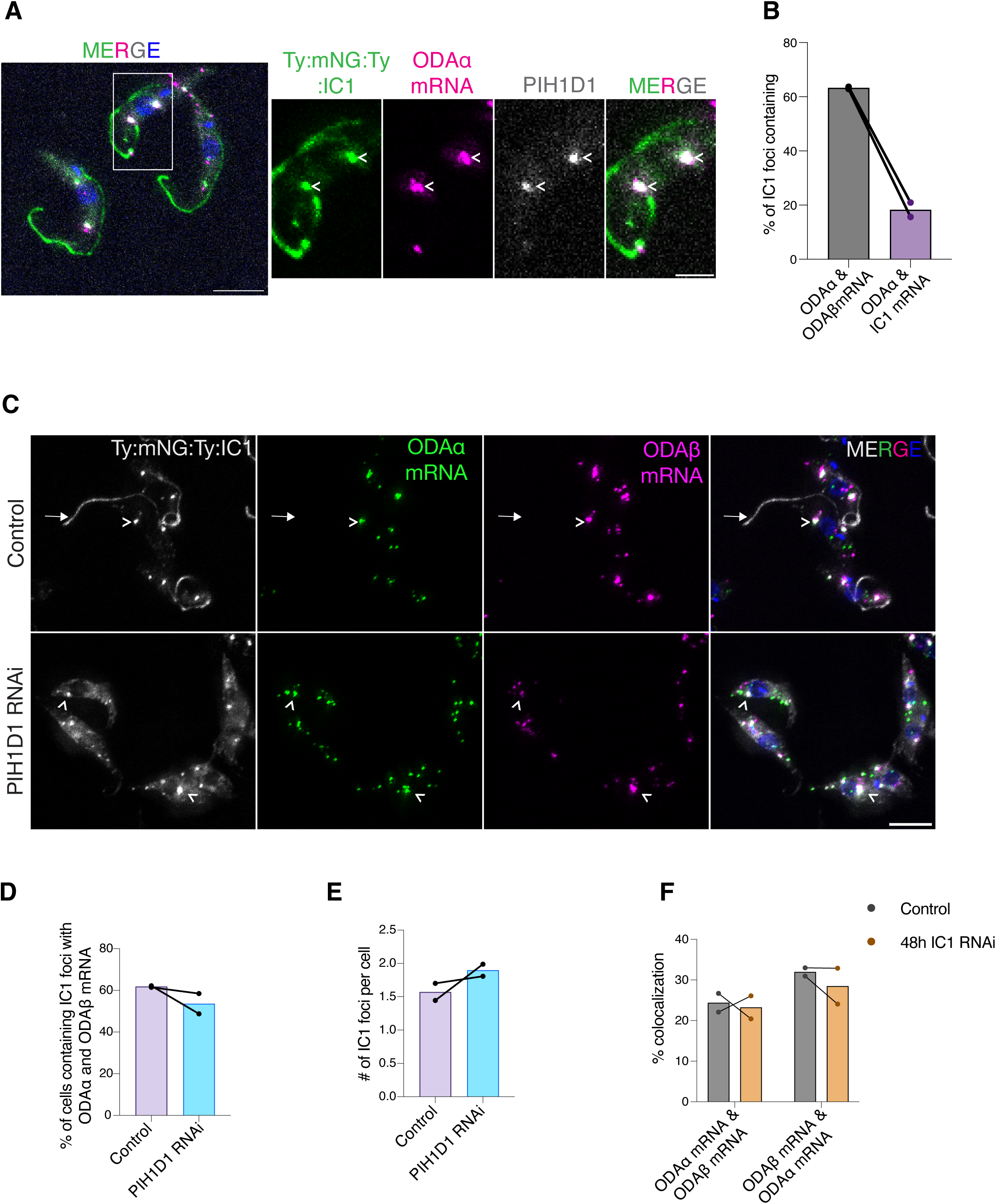
IC1 is post-translationally recruited to the ODA HC translational sites. **A)** Representative images of Ty:mNG:Ty:IC1 cells co-labelled for ODAα mRNA and PIH1D1. Arrowheads indicate the colocalization of IC1 protein with ODAα mRNA and PIH1D1. Scale bar = 5 µm; Inset scale = 2 µm. **B)** Quantitation plot showing % IC1 foci containing both ODAα mRNA and ODAβ mRNA or both ODAα mRNA and IC1 mRNA. The lines represent the number of biological replicates. **C)** Ty:mNG:Ty:IC1 cells were induced for PIH1D1 RNAi (48h) or not, and labelled for ODAα mRNA and ODAβ mRNA. Arrows indicate the flagellum. Arrowheads point to the colocalization of IC1 protein with both ODAα mRNA and ODAβ mRNA. Scale bar = 5 µm. **(D-E).** Quantitation plot displaying % of cells containing IC1 foci with ODAα mRNA and ODAβ mRNA or number of IC1 foci per cell, with or without PIH1D1 RNAi (48h). At least 200 cells were analyzed per biological replicate. **F)** The effect of IC1 RNAi (48h) on the colocalization of ODAα mRNA and ODAβ mRNA. The effect was quantitated as described in Fig. 6A and 6B. For each biological replicate, at least 500 cells were analyzed. The lines in the graph represent the number of biological repeats.

We also examined how IC1- and ODAβ-RNAi affect ODAα translation using the MoonTag-ODAα reporter cell line. Surprisingly, though the HCs depend on each other for stability (Tingting, 2022), ODAβ RNAi did not affect the number of ODAα translational sites per cell (Fig. 6F). On the contrary, IC1 RNAi significantly reduced the number of ODAα translational sites per cell (Fig. 6F). Additional smFISH experiments showed that IC1 is not required for bringing the mRNA of the two heavy chains together, and ODAα and ODAβ colocalized regardless of IC1 depletion (Fig. 7F). Thus, IC1 is likely recruited after the initial HC assembly and is essential to synthesize HCs efficiently.

## Discussion

In this study, we characterized the molecular landscape of a conserved subset of DNAAFs known as the PIH family proteins in *Trypanosoma brucei*. We identified PIH1D1 as an essential DNAAF for ODA assembly (Fig. 2, Fig. S2, Fig. 3, Fig. 6). Intriguingly, PIH1D1 forms cytoplasmic foci that contain ODA subunit IC1 as well as translating heavy chains (Fig. 4, Fig. 5, Fig. 7), and are more abundantly present in cells building a new flagellum (Fig. 4B & S4B). The PIH1D1 foci are reminiscent of DynAP-like compartments reported in other eukaryotes (see below). This study provides direct evidence supporting a co-post-translational assembly model for the ODA (Fig. 8), lending insights into the spatiotemporal organization and regulation of ODA biogenesis.

**Figure 8.**
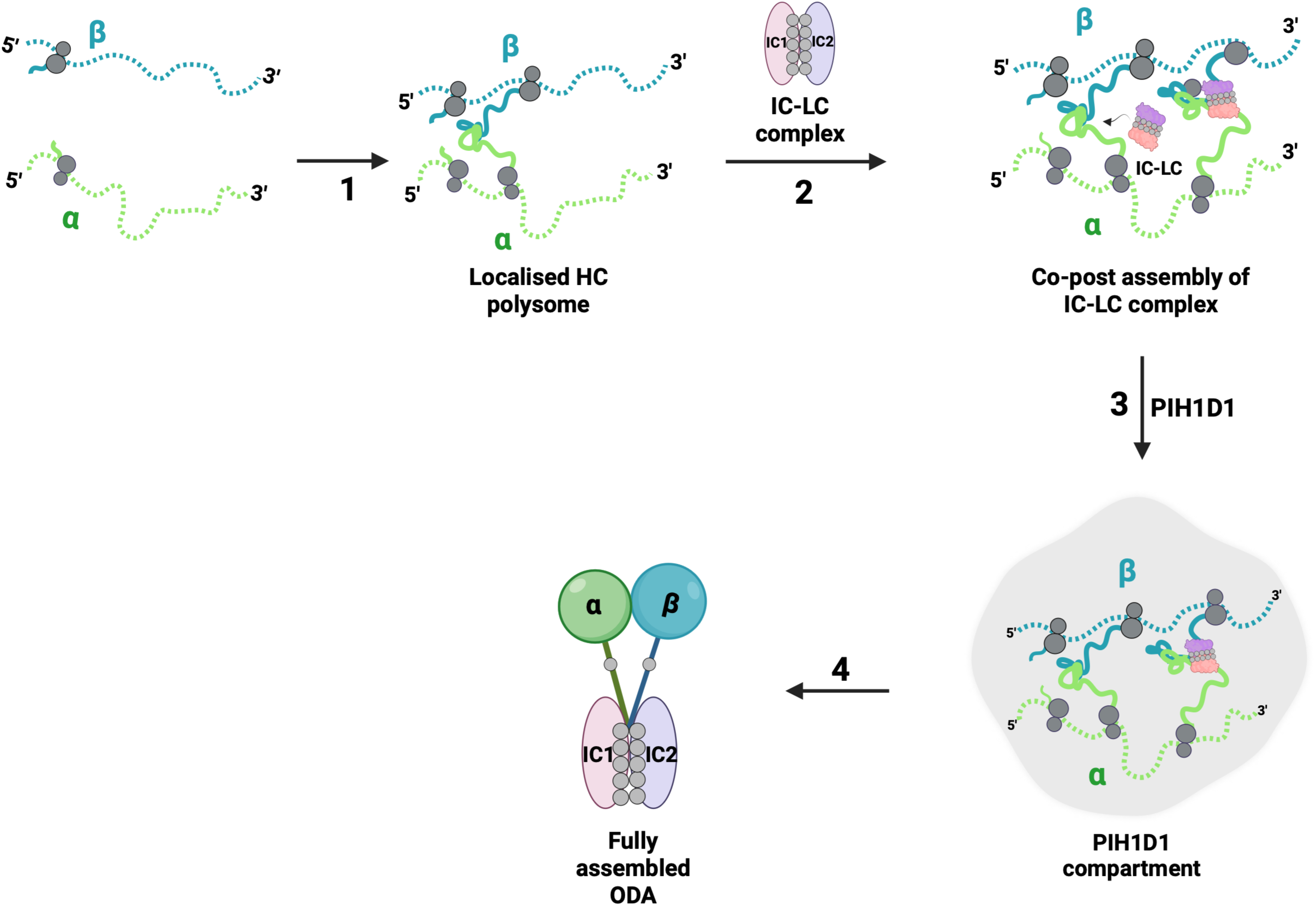
Spatio-temporal regulation of ODA. Schematic diagram depicting the order of events leading to the full ODA formation. 1. The initial co-translational pairing of the heavy chains ODAα and ODAβ. 2. The IC-LC complex is post-translationally recruited to assemble with the HCs as they are being synthesized. 3. & 4. Following the IC-LC recruitment, PIH1D1 forms specialised compartments at the co-translational assembly sites to promote the folding of HCs and the stable formation of ODA (Created in BioRender. Balasubramaniam, K. (2025) https://BioRender.com/4s75e3y).

How the ODA subunits navigate and find each other in the cytosol to form a functional ODA with fixed stoichiometry has been a long-standing question. Of the ODA subunits, the HCs are the largest polypeptides, each made up of ∼4600 amino acids. The N-terminal region of the HCs encodes an N-terminal dimerization domain (NDD) that mediates direct interaction between ODAα and ODAβ (Mali et al., 2021; Walton et al., 2021). The NDDs are followed by the domains required for interaction with the IC-LC complex and other LC subunits such as LC1. Here we demonstrated that ODAα and ODAβ dimerize co-translationally, likely via interaction between their NDDs. This is followed by the deposition of preassembled IC-/LC complex onto the elongating heavy chains (Fig. 8). These findings are supportive of a hypothetical model proposed by Stephen King, where the IC, LC subunit association is coupled to HC translation (King, 2021). While the ICs are not necessary for the cotranslational dimerization of the HCs, they play a critical role in efficiently translating HCs. It remains to be seen how IC deficiency, therefore the lack of IC-LC complex, might affect HC translation elongation, and how the IC-LC complex itself is assembled and regulated. Our study highlights a critical role of ICs as an assembly checkpoint, to ensure productive HCs synthesis only in the presence of the partner IC-LC complex and thus correct stoichiometry of the final assembled products.

Importantly, our results demonstrate a function of DNAAF, PIH1D1, at a late step in ODA assembly, downstream of IC-LC association with translating HCs. First, the depletion of PIH1D1 did not affect the co-translational association of ODAα and ODAβ, nor the enrichment of the IC-LC complex at the HC co-translational sites. Second, the core translational steps, such as the initiation or elongation of the heavy chains, are also independent of PIH1D1. Finally and importantly, the localization of PIH1D1 to discrete cytoplasmic foci is dependent on the expression of both HCs and IC1. Based on these findings, we posit a model where the assembly of IC-LC complex with translating HCs further nucleates the formation of PIH1D1 foci (Fig. 8). Given the abundance of chaperones and assembly factors identified in PIH1D1-BioID, we propose that PIH1D1 acts by concentrating relevant chaperones and assembly factors at the cotranslational assembly sites of ODA to facilitate proper folding and stabilization of assembly intermediates. In the absence of PIH1D1, the newly synthesised HCs may not be properly folded and cannot form a stable assembly with the IC-LC complex, resulting in rapid degradation.

Chaperones engaging with nascent polypeptides is a conserved strategy to shield nascent proteins from premature degradation or spurious interactions. Examples of such chaperones in eukaryotes include Ssb of the Hsp70 family (Willmund et al., 2013) and the Nascent Chain Associated Complex (NAC) (del Alamo et al., 2011). Recently, a study (Philippe et al., 2025) has shown that the components of the co-chaperone R2TP, such as RUVB1 and RPAP3, bind to several clients not only post-translationally but also co-translationally. The study also suggested HSP90/R2TP forms chaperone factories to facilitate the co-translational interactions with their client proteins. Similarly, PIH1D1 may form chaperone factories to aid specific steps during the cotranslational assembly of ODA. In addition to PIH1D1, the other PIH homologs KTU, PIH1D2, and PIH1D3 may also act co-translationally on their respective clients.

Many DNAAFs have been reported to form discrete cytoplasmic foci in higher eukaryotes (Diggle et al., 2014; Horani et al., 2018; Y. Li et al., 2017). But, the requirement for a dedicated membraneless organelle for axonemal dynein assembly was first reported in *Xenopus* (Huizar et al., 2018; Lee et al., 2020). These organelles, called DynAPs (Dynein Axonemal Assembly Particles), are dynamic, liquid-like condensates that concentrate axonemal dynein subunits along with multiple DNAAFs, including PIH proteins and molecular chaperones. Although RNA has been detected in these organelles (Drew et al., 2020), their identity is not known, and the exact function of DynAPs is still speculative. The functions of DynAP-like organelles were studied in *Drosophila* (Fingerhut & Yamashita, 2020) and, more recently, in *Zebrafish* (Li et al., 2024). In *Drosophila*, these organelles, termed “Kl granules,” appear during spermatogenesis to cluster ODA heavy chains mRNA but not their proteins, suggesting a role as mRNA storage sites. In *Zebrafish,* DynAP-like structures contain translating ODA heavy chains, supporting their role as sites of co-translational dynein heavy chain assembly. Here we propose that the PIH1D1 compartments in *T. brucei* are DynAP-like organelles, serving as late-stage cotranslational ODA assembly sites. Although there may be organism-or cell-specific differences, it appears that spatial compartmentalization is a conserved strategy for ordered ODA assembly from protists to humans.

Overall, our study reveals that the assembly of ODA in the cytosol is highly organized with multiple layers of assembly checkpoints. By spatially confining the HCs, the cells coordinate the translation of the HCs to their orderly interactions with other ODA subunits. Once the translation of the HCs is complete, the fully assembled ODA is released and trafficked to the cilia (Fig. 8).

## Materials and Methods

### Cell lines and maintenance

The procyclic form of *Trypanosoma brucei,* strain 29.13 or DIY (F. J. Li et al., 2017; Wirtz et al., 1999), was used to generate all the cell lines in this study. These cells were cultured in Cunningham medium supplemented with 10% heat-inactivated fetal bovine serum (FBS) (Hyclone), appropriate antibiotics, and incubated at 28°C. The 29.13 strain engineered for tetracycline-inducible expression was used to create PIH1D1 RNAi cell lines. All cell lines expressing endogenously tagged mNG or YFP (ODAα, ODAβ, IC1, and IC2) were derived from 29.13 cells containing the PIH1D1 RNAi inducible system. To express cumate-inducible BioID2 tagged-PIH homologs (N-term), the DIY strain engineered for dual induction with cumate and tetracycline was employed. All MoonTag-related cell lines were generated using the DIY strain.

### Plasmid construction

All the gene sequences were downloaded from Tritryp Database (https://tritrypdb.org/tritrypdb/app) (Alvarez-Jarreta et al., 2024) (Shanmugasundram et al., 2023). All plasmids were constructed using the traditional or Gibson assembly-based cloning techniques. The gene fragments targeted for RNAi were selected using the RNAit server (https://dag.compbio.dundee.ac.uk/RNAit/) (Redmond et al., 2003). Detailed information on the plasmids used in this study is shown in Table S1.

### Cell growth assay

The cells were inoculated at a cell density of 0.5×10^6^ cells/ml and monitored every 24h. If the cell density exceeded 1×10^6^ cells/ml, the cells were diluted to either 0.5×10^6^ cells/ml or 1×10^6^ cells/ml. The doubling number was calculated using the formula: Log^2^[Df*(N^t^/N^0^)]. Df is the cumulative dilution factor, N^t^ is the cell density at a given time point (t), and N^0^ is the cell density at time=0.

### Detergent extraction

A total of 2×10^6^ cells were harvested by centrifugation at 800 g for 5 min and washed twice with ice-cold 1X PBS supplemented with EDTA-free protease inhibitors (Roche). The cells were resuspended in 50 µl of PEME buffer (0.1 M PIPES pH 6.9, 2 mM EGTA, 1 mM MgSO4 and 0.1 mM EDTA) containing 1% NP-40 and protease inhibitors (Roche), and extracted on ice for 5 min. Extracted cells were centrifuged at 20,000 g for 10 min at 4°C. The detergent soluble supernatant was transferred to a fresh tube, and the detergent-insoluble pellet was resuspended in 50 µl of PEME buffer. 50 µl of 2X SDS loading buffer was added to both detergent-soluble and -insoluble fractions and boiled for 5 min. The boiled protein samples were stored at −80⁰C or used immediately for immunoblotting analyses. To obtain total cell lysates, 2×10^6^ cells were harvested, washed, and extracted as above. 50 µl 2X SDS loading buffer was added directly to the total cell lysates, and the samples were boiled. This protocol was adapted from (Alves et al., 2020).

### Immunofluorescence assays and imaging

For whole-cell fixation, *T. brucei* cells were harvested at 1800 g for 1 min, washed twice with 1X PBS, and attached to the cover glass for 10 min. The attached cells were fixed with 4 % PFA and permeabilized with 0.25 % Triton-X 100 (in 1X PBS) for 10 min each. The permeabilized cells were either incubated with chilled methanol (−20⁰C) for 5 min and rehydrated in 1X PBS for 10 min, or not, before blocking with 3 % BSA. For *in situ* fixation, PFA was added directly to the cell culture to a final concentration of 4% and incubated for 15 min at RT with gentle shaking. The cells were then harvested at 1800g for 1 min and washed twice with 1X PBS and sedimented onto the cover glass by centrifugation. The cells were then permeabilized with 0.25 % Triton-X 100 (in 1X PBS), blocked with 3% BSA, and then stained with respective antibodies (primary & secondary) and DAPI (25 µg/ml) following the conventional protocol. The antibodies used in this study are listed in Table S2.

For the translation inhibition assays, the cells were treated with puromycin (200 µg/ml) for 30 min or harringtonine (3 µg/ml) for a specified time. Drug-treated cells were fixed *in situ* with 4 % PFA for 15 min and quenched with 30 mM glycine for 5 min, both at room temperature. After quenching, the samples were placed on ice or stored at 4⁰C until further processing. All images were acquired with a 63x-1.4 NA objective and a Cool SNAP HQ2 CCD camera (Photometrics) on a Zeiss Axio Observer Z1 fluorescence microscope.

### Image processing and statistical analysis

All image processing and quantitation were carried out using the plugins available on Fiji. For the flagellar intensity measurement, the anterior tip of the flagellum (∼1 µm) was measured using a specified line length tool. For image display, the z-stack images were projected using the Max Intensity Projection type. For the smFISH experiments, ComDet v.0.5.6 plugin was used to count and assess the colocalization of the mRNA particles, PIH1D1 foci, and MoonTag-ODAα translational sites. The maximum distance between the colocalized spots was set to 4 pixels. The approximate particle size and the intensity threshold were adjusted manually to include all the distinct particles. The Z-stack images were analysed directly or processed with maximum intensity projection type before the ComDet analysis. The Dot Plot, Venn Diagram and heatmaps were generated using R (version 4.4.1).

GraphPad Prism 9 (Version 9.5.1 (528)) was used to prepare all the graphs and perform statistical tests. Unpaired t test with Welch’s correction was performed. *p* <0.05 was considered statistically significant.

### Purification of cytosolic ODA complex

For sucrose gradient analysis, the cytoplasmic extracts were prepared by treating the cells with PEME buffer (0.1 M PIPES, pH 6.9, 2 mM EGTA, 1 mM MgSO4, 0.1 mM EDTA) containing 1% NP-40 and protease inhibitors (Roche) for 10 min on ice. The cell lysates were then centrifuged at 20,000 g for 10 min at 4°C. The supernatant was gently placed on top of a 5-30% sucrose gradient (12 ml) and ultracentrifuged for 16 h at 36,000 rpm using the SW41 rotor. 0.5 ml fractions were collected from the top of the gradient manually and processed for immunoblot analysis.

For affinity purification, PIH1D1 RNAi cells expressing endogenously tagged Ty:YFP: Ty:IC1 were collected with or without RNAi induction after 72 h. Cells expressing GFP-Ty only were used as a control. Briefly, 5×10^8^ cells were harvested at 3000 rpm for 7 min at room temperature. The cell pellets were washed with ice-cold polysome buffer (20 mM Tris-HCl pH 7.5, 120 mM KCl, 5 mM MgCl2, 1 mM DTT, and 200 mM sucrose, supplemented with Protease Inhibitor Cocktail (Roche) (Bajak & Clayton, 2020). In Trial 1, 100 µg/ml cycloheximide was also included in the wash buffer. The cell pellet was then flash frozen in liquid nitrogen and stored at −80⁰C until further processing. The frozen cell pellets were resuspended gently and thoroughly in 350 µl of lysis buffer (1 % Triton-X 100, 20 mM Tris-HCl pH 7.5, 120 mM KCl, 5 mM MgCl2, 1mM DTT, 200 mM sucrose), containing 100 µg/ml cycloheximide or not. The cell suspension was passed 10 times through 25 ½ and 27 ½ 1 ml syringe needle consecutively and clarified by centrifuging at 15000 g for 10 min at 4⁰C. The clarified cell lysate was diluted with 300 µl of dilution buffer (20 mM Tris HCl pH 7.5, 120 mM KCl, and 5 mM MgCl2, supplemented with Protease Inhibitor Cocktail) and incubated with 50 µl of equilibrated GFP-Trap® Magnetic Agarose beads (ChromoTek) for 1 h on a rotator at 4⁰C. The GFP-bound magnetic agarose beads were washed thrice with lysis buffer (without cycloheximide) and once with 1X PBS. 200 µl triethylammonium (500 mM, pH 8.5) and 4 µl 100 mM TCEP were added to the beads and incubated for 1 h at 50⁰C in a rotary shaker. To alkylate the proteins, the beads were further incubated with 5 mM MMTS for 15 min at room temperature. Lastly, the bead-bound proteins were digested with 2.5 µg of Trypsin overnight at 37⁰C in a rotary shaker, and desalted for mass spectrometry analysis.

### Proximity-dependent biotinylation using BioID2

3xHA-BioID2 fusion to all four PIH homologs was expressed using a cumate-inducible system. After induction with 10 µg/ml cumate for 8 h, cells were incubated with 50 µM biotin for an additional 16 h. Wild-type cells without BioID2 expression, ∼1-2×10^9^ cells, were harvested at 3000 rpm for 7 min (RT) and washed thrice with 1X PBS. The pelleted cells were lysed thoroughly in 1% Triton X-100 (prepared in 1XPBS containing Protease Inhibitor Cocktail) and the cell lysates were clarified by centrifuging at maximum speed for 10 min at 4⁰C. The clarified cell lysates were incubated with 50 µl of streptavidin beads (Dynabeads® M-280 Streptavidin) for 4 h in a rotary shaker at 4⁰C. The proteins bound to streptavidin beads were separated using a magnetic stand and washed stringently once with 1% SDS in 1XPBS, twice with 1% Triton X-100 in 1XPBS, and thrice with 1XPBS for 5 min for each wash. All washes were performed at room temperature. After washing, the protein-bound streptavidin beads were separated and subjected to on-bead digestion as described previously.

### Fluorescent *in situ* RNA hybridization using View RNA ISH cell plus assay

3-4 x 10^6^ cells/ml were collected and fixed *in situ* as described above. The cells were washed twice with 1X PBS (Supplemented with RNase inhibitor) and attached to 5 mm round cover glass in a 96-well plate by centrifugation at 1800g for 30 min. Samples were processed for simultaneous detection of protein and mRNA using the ViewRNA® Cell Plus Assay kit (Catalog No:188-19000-99, Thermo Fisher Scientific). The experiment was performed following the manufacturer’s protocol optimized for a 96-well plate format with some modifications. Custom proprietary RNA probe sets for detecting ODAα (Tb927.3.930, Type 1 probe, Assay ID-VPRWENM), ODAβ (Tb927.11.3250, Type 4 probe, Assay ID:VPU62XJ or Type 6 probe, Assay ID:VPWCWHG), IC1 (Tb927.11.5000, Type 6 probe, Assay ID: VPPRJ6F) were designed by the manufacturer based on full length coding sequences.

### RNA extraction and qRT-PCR

Total RNA was isolated from a total of 3-4×10^7^ cells using FreeZol reagent (R711-01, Vazyme) according to the Manufacturer’s manual. 1 µg of RNA was used to synthesize cDNA using the HiScript III RT SuperMix for qPCR (+gDNA wiper) Kit (R323-01, Vazyme). For qPCR analysis, the reactions were prepared using Taq Pro Universal SYBR qPCR Master Mix (Q712-02, Vazyme) and analysed on the CFX96 Touch Real-time system (Bio-Rad, California, USA). The relative expression of all gene targets was normalised against TERT (Brenndorfer & Boshart, 2010). Three independent biological replicates were performed, each with three technical replicates. The primers used for the analysis are listed in Table S3.

## Supplementary Figure Legends

**Figure S1.**
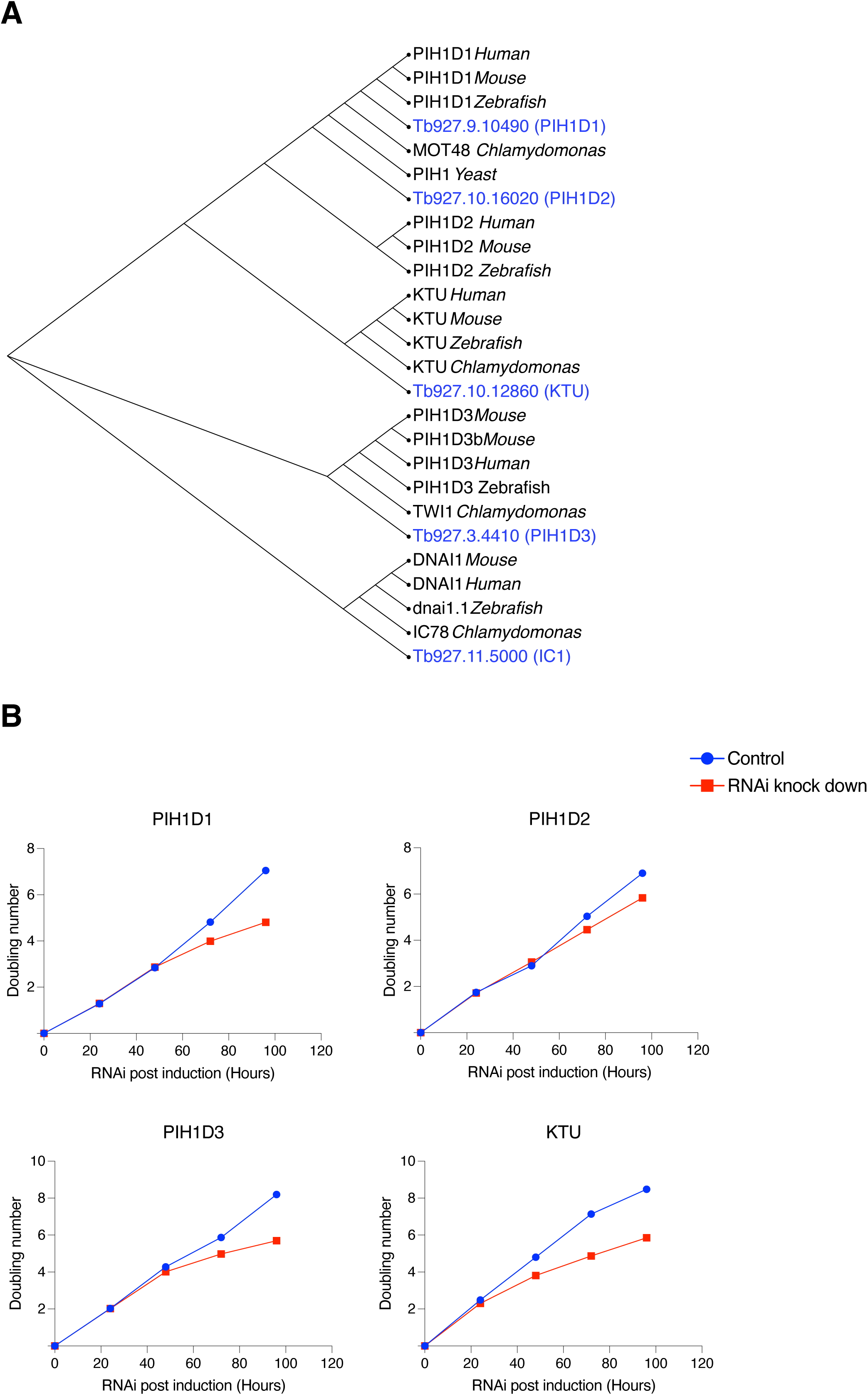
PIH homologs are crucial to cell growth and motility. **A)** Phylogenetic tree of PIH homologs. Multiple sequence alignment was performed using Clustal Omega, and a phylogenetic tree was generated by the neighbor-joining method with default parameters available on the EMBL-EBI website https://www.ebi.ac.uk/. The tree obtained was annotated using iTOL (https://itol.embl.de/). ODA intermediate chain 1 (DNAI1 or IC1) was used as a reference group. **B)** Growth curve analysis post RNAi induction of all four PIH homologs.

**Figure S2.**
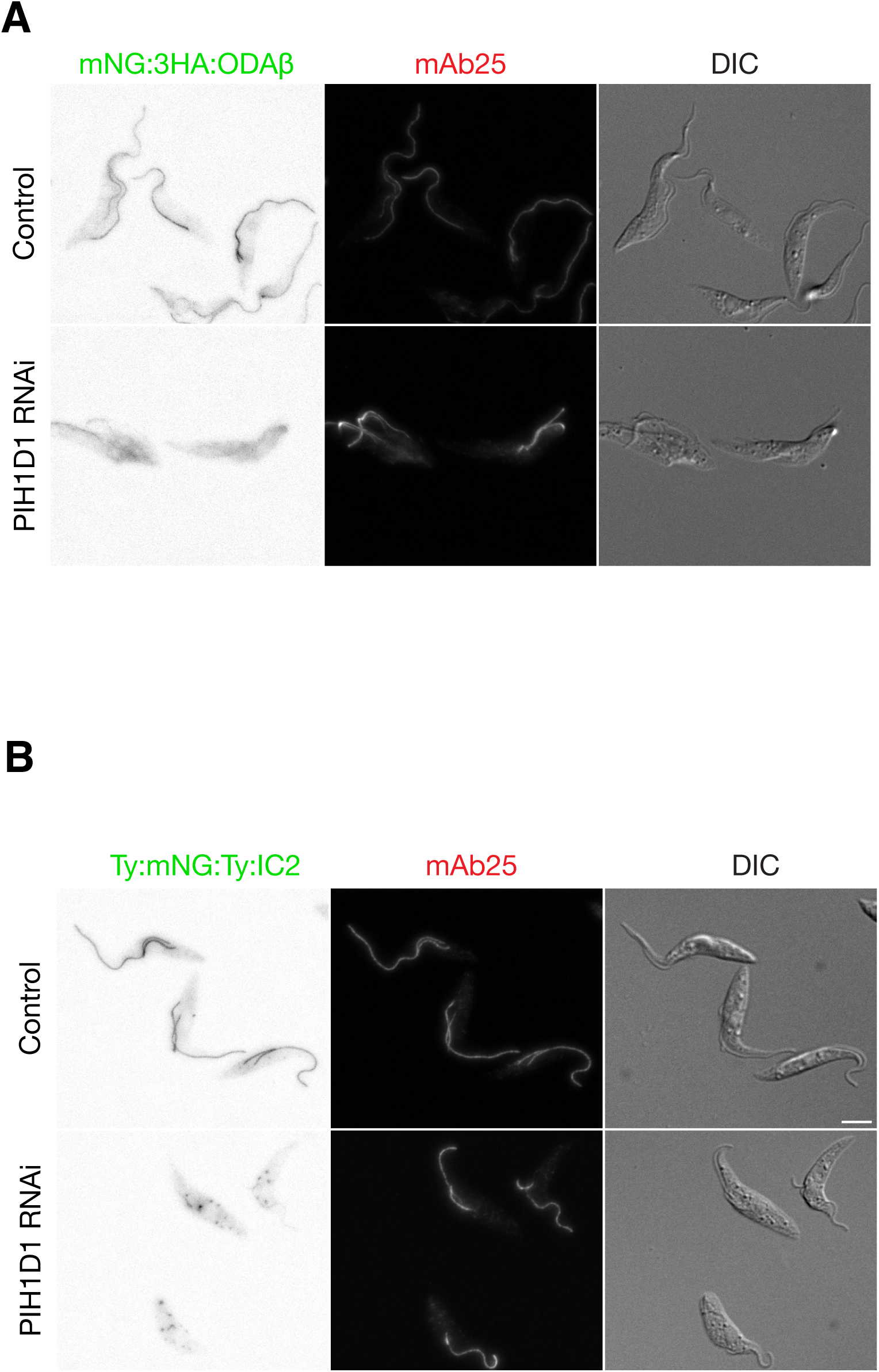
Depletion of PIH1D1 reduces the protein abundance of ODA HCs but not the ICs. **(A-B)** Cells endogenously expressing ODAβ or IC2 tagged N terminally with mNG:3HA or Ty:mNG:Ty were induced for PIH1D1 RNAi (72h). Uninduced cells were used as controls. The cells were co-labelled with mAb25 to mark the axoneme. Scale bar = 5 µm.

**Figure S3.**
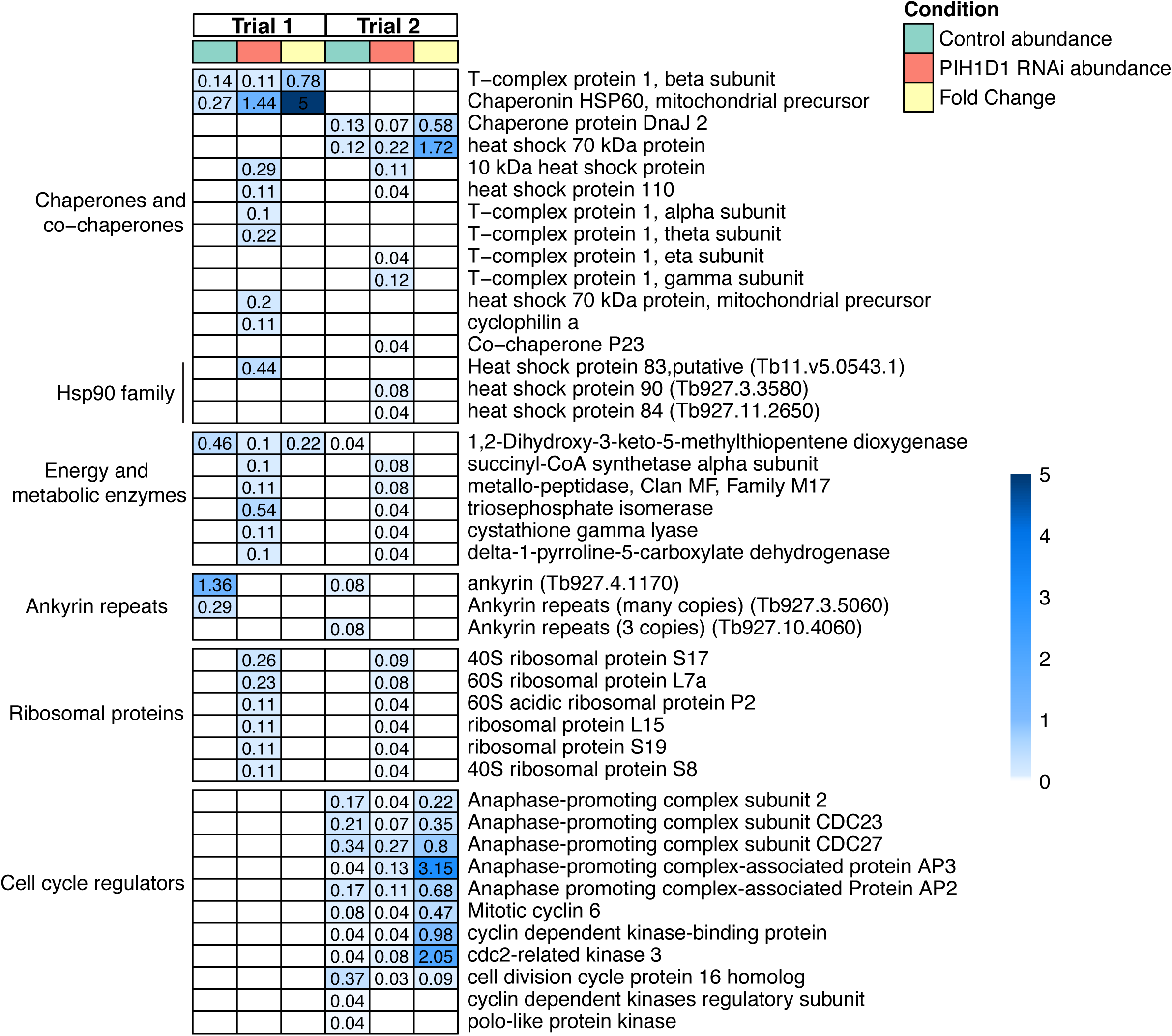
List of non-ODA and cytoplasmic proteins co-purified with IC1. Heatmap showing the protein abundance and fold change of IC1-associated proteins representing different functional categories, identified in control and PIH1D1 RNAi (72h) cells.

**Figure S4.**
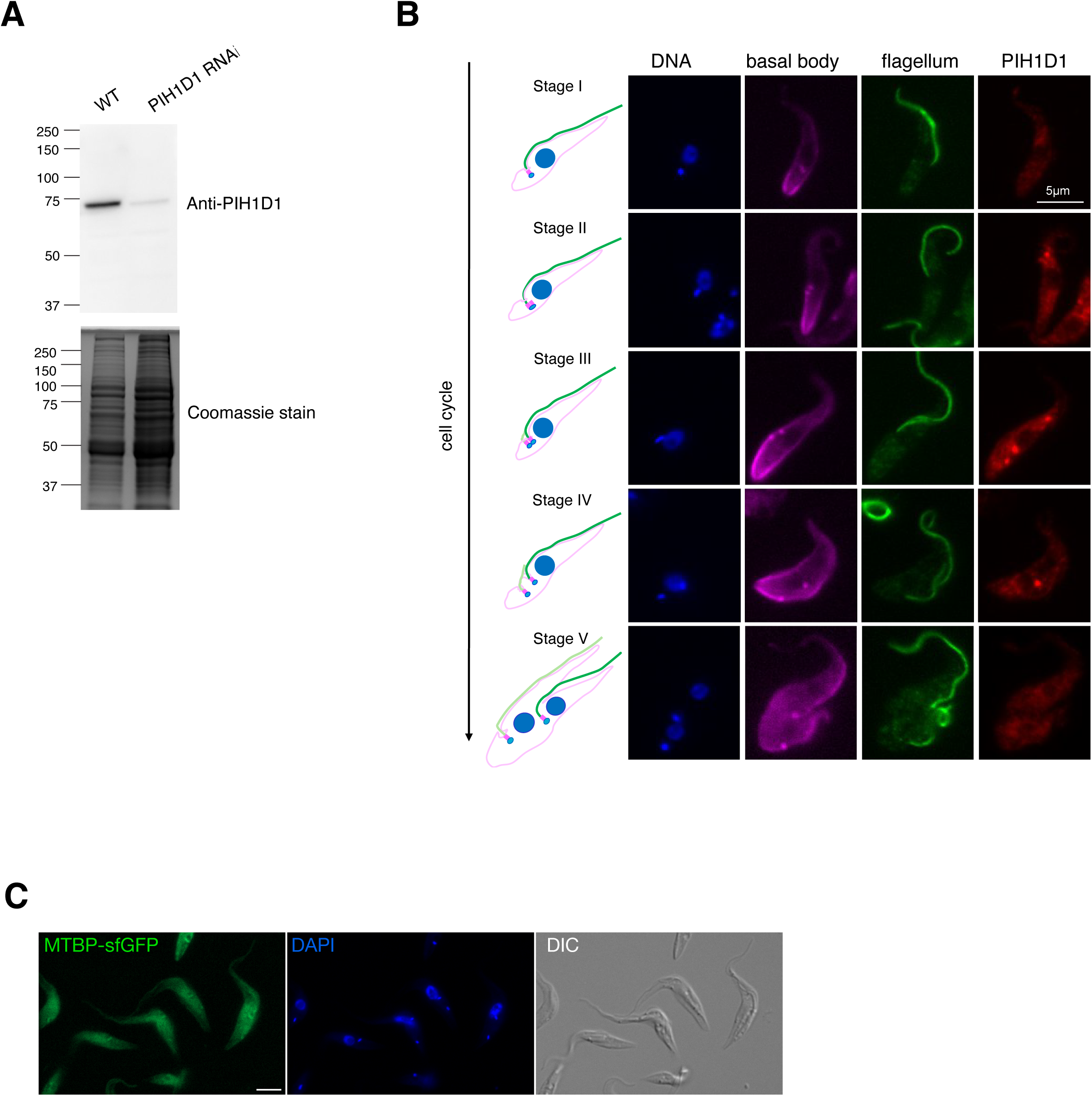
PIH1D1 foci formation across different cell cycle stages. **A)** Validation of the anti-PIH1D1 antibody by immunoblot analysis. Strong, specific labelling of a ∼72 KDa band with size expected for the native PIH1D1 protein was detected in the wild-type cells (29.13) but weakly in the cells induced for PIH1D1 RNAi (72 h). Total proteins stained with Coomassie brilliant blue were used as a loading control. **B)** Cells were immunolabelled with anti-PFR that marks the flagellum, YL1/2 antibodies that mark the basal body, and a polyclonal anti-PIH1D1 antibody to track the PIH1D1 puncta formation across different cell cycle stages. DNA was visualised using DAPI. Construction of a new flagellum occurs soon after duplication of the basal bodies. As the new flagellum continues to elongate, the duplicated basal bodies drive the division of the duplicated kinetoplast (mitochondrial DNA, small dots). Once the nuclear division completes, the biflagellated mother cells enter cytokinesis. One representative cell in different cell cycle stages is shown along with the cartoon depiction. Scale bar = 5 µm. **C)** Cells constitutively expressing MoonTag binding protein fused to sfGFP (MT-sfGFP) display a diffuse cytoplasmic signal. DNA is stained with DAPI. Scale bar = 5 µm.

## Supplementary Data

**Data 1.** List of PIH homologs BioID hits.

**Data 2.** Mass spectrometry analysis of affinity-purified YFP-IC1 samples from control and PIH1D1 RNAi (72h).

## Supplementary Tables

**Table S1.**
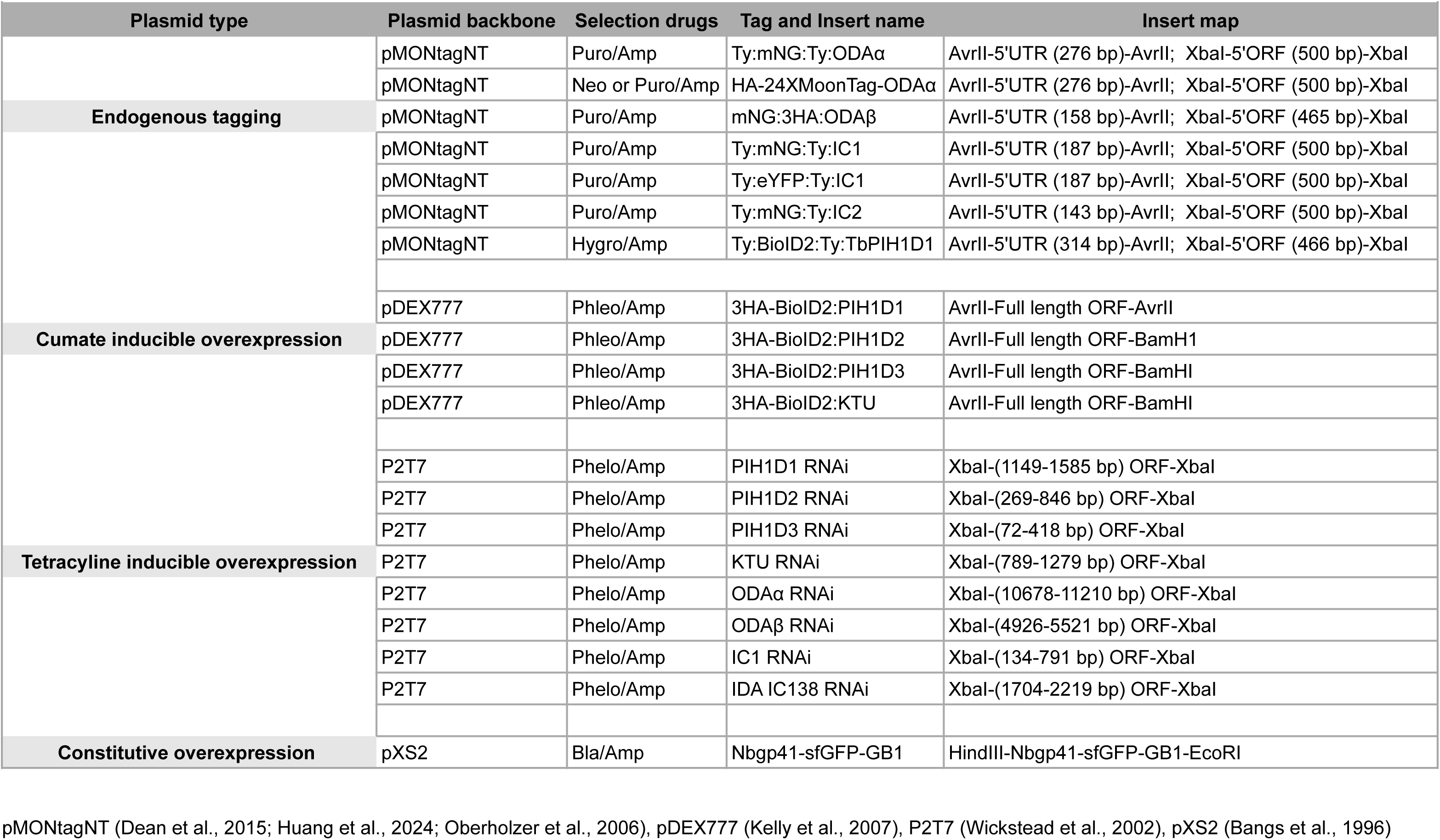
List of plasmids used in this study.

**Table S2.**
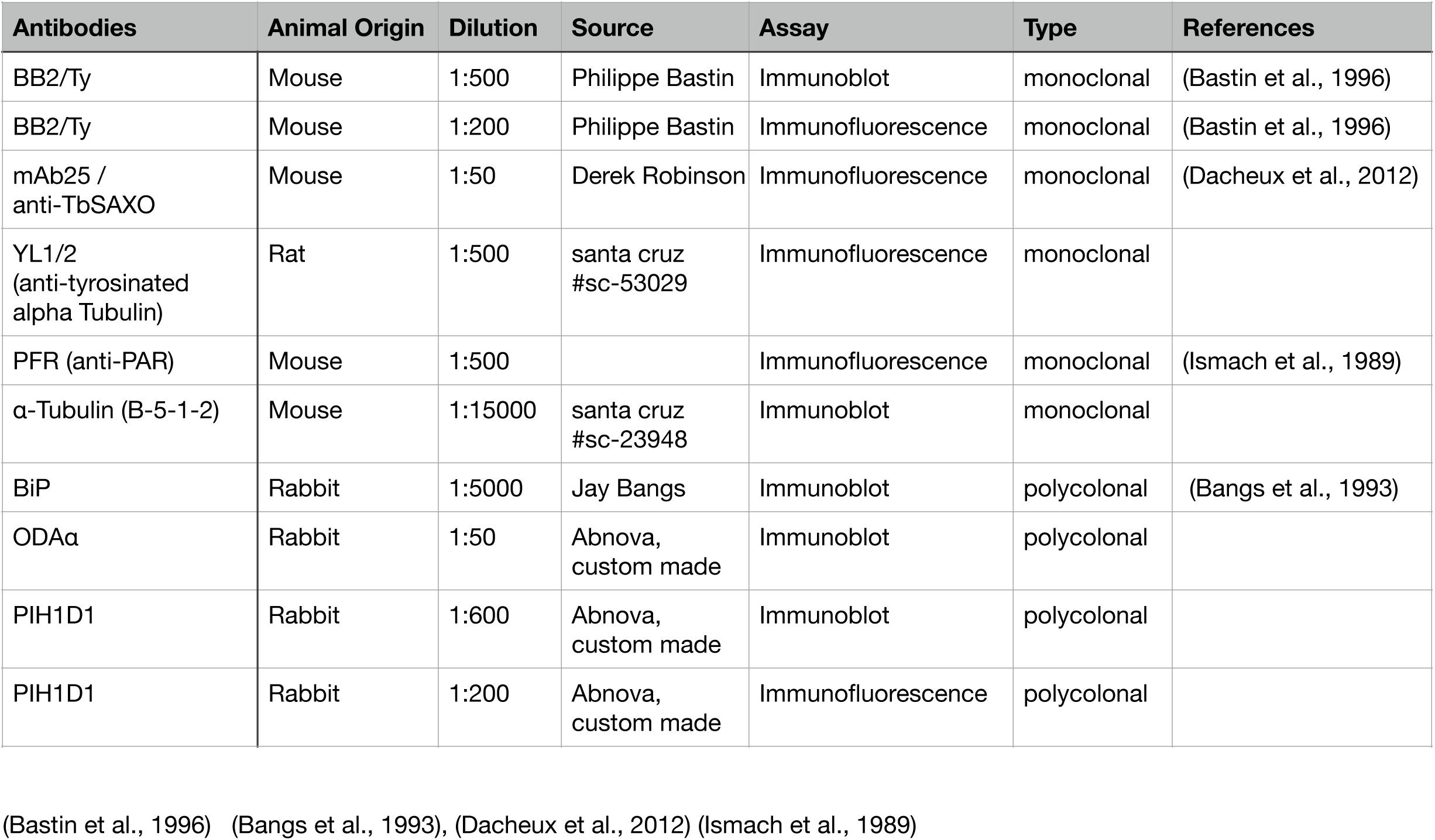
List of antibodies used in this study.

**Table S3.**
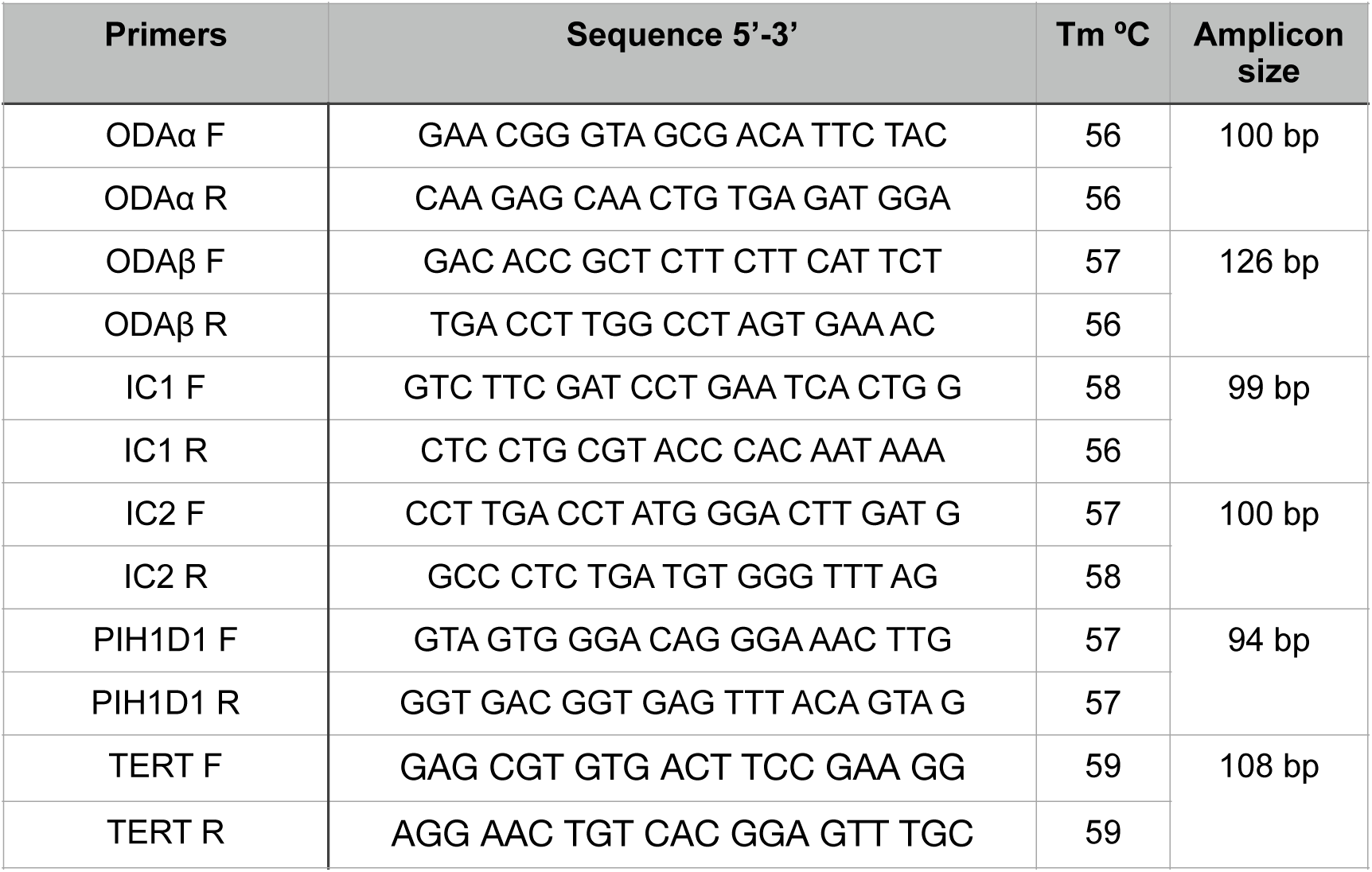
List of primers used for qRT-PCR analysis.

## Author contributions

K.B. developed the hypothesis and designed the study with input from C.Y.H. K.B. performed most of the experiments. F.F. purified PIH1D1 for antibody generation, validated the antibodies, and performed cell-cycle analyses of PIH1D1 foci. K.B. and C.Y.H. analyzed and validated the data. C.Y.H. supervised the project. K.B. wrote the original draft. C.Y.H. and K.B. revised and edited the draft.

## Acknowledgements

We thank Qingsong Lin, Teck-Kwang Lim, and Xin Ee Yong at the Protein and Proteomics Centre (PPC), NUS, for mass spectrometry analysis; Tingting He for providing technical assistance; Helen Chen for generating the anti-ODAα antibodies; Helen Chen and Darren Teo for sucorse gradient experiment; Hao Yuan Yang for initial work on optimizing constructs for expressing translational reporters in *T. brucei*; Dr. Zhewang Lin, Dr. Shifeng Xue, and Dr. Bor Leun Tang for fruitful discussions; and Dr. Zhewang Lin for providing harringtonine reagent and advice on the experiment. This work was supported by the Singapore Ministry of Education Tier 2 research grant MOE-T2EP30121-0003 awarded to C.Y.H. K.B. was a recipient of the NUS research scholarship.

## Competing interests

The authors declare no competing interests.

## Data and materials availability

All data needed to evaluate the conclusions in the paper are present in the paper and/or the Supplementary Materials. The proteomics results will be deposited to PRIDE.

## Notes

### Competing Interest Statement

The authors have declared no competing interest.

